# Detection of shared balancing selection in the absence of trans-species polymorphism

**DOI:** 10.1101/320390

**Authors:** Xiaoheng Cheng, Michael DeGiorgio

**Affiliations:** Huck Institutes of the Life Sciences, Pennsylvania State University, University Park, PA, USA; Department of Biology, the Pennsylvania State University, University Park, PA, USA; Department of Statistics, the Pennsylvania State University, University Park, PA, USA; Institute for CyberScience, the Pennsylvania State University, University Park, PA, USA

## Abstract

Trans-species polymorphism has been widely used as a key sign of long-term balancing selection across multiple species. However, such sites are often rare in the genome, and could result from mutational processes or technical artifacts. Few methods are yet available to specifically detect footprints of trans-species balancing selection without using trans-species polymorphic sites. In this study, we develop summary- and model-based approaches that are each specifically tailored to uncover regions of long-term balancing selection shared by a set of species by using genomic patterns of intra-specific polymorphism and inter-specific fixed differences. We demonstrate that our trans-species statistics have substantially higher power than single-species approaches to detect footprints of trans-species balancing selection, and are robust to those that do not affect all tested species. We further apply our model-based methods to human and chimpanzee whole genome sequencing data. In addition to the previously-established MHC and malaria resistance-associated *FREM3/GYPE* regions, we also find outstanding genomic regions involved in barrier integrity and innate immunity, such as the *GRIK1/CLDN17* intergenic region, and the *SLC35F1* and *ABCA13* genes. Our findings not only echo the significance of pathogen defense, but also reveal novel candidates in maintaining balanced polymorphisms across human and chimpanzee lineages. Finally, we show that these trans-species statistics can be applied to and work well for an arbitrary number of species, and integrate them into open-source software packages for ease of use by the scientific community.

## Introduction

Balancing selection is an evolutionary mechanism for maintaining diversity within populations (Charlesworth, 2006). A number of different modes of balancing selection exist, such as heterozygote advantage (Charlesworth, 2006), pleiotropy (Johnston et al., 2013), negative frequency-dependent selection (Mitchell-Olds et al., 2007), environmental fluctuations (Bergland et al., 2014), and segregation distortion balanced by negative selection (Charlesworth and Charlesworth, 2010; Ubeda and Haig, 2004). Though these different modes vary in how they maintain polymorphism over long periods of time, they all leave behind similar genomic signatures of increased density of polymorphic sites nearby a balanced polymorphism, and often an enrichment of middle-frequency alleles in a narrow window surrounding the selected locus (Charlesworth, 2006). These characteristic footprints have been utilized by a number of statistical approaches for detecting long-term balancing selection (*e.g*., Hudson et al., 1987; Tajima, 1989; Andrés et al., 2009; Leffler et al., 2013; DeGiorgio et al., 2014; Gao et al., 2015; Hunter-Zinck and Clark, 2015; Sheehan and Song, 2016; Siewert and Voight, 2017; Bitarello et al., 2018).

However, until recently, the availability of methods for detecting long-term balancing selection has been limited, and the most commonly-used approaches were the Hudson-Kreitman-Aguadé (HKA; Hudson et al., 1987) and Tajima’s *D* (Tajima, 1989) statistics. While the HKA statistic captures increases in polymorphism density, it does not consider allele frequency information, and therefore cannot sense the enrichment of intermediate-frequency alleles. On the other hand, Tajima’s *D* measures the distortion of the allele frequency spectrum from the expectation under a constant size neutrally-evolving population, and has the ability to identify the footprint of an increased number of middle-frequency alleles in a genomic region. However, as it does not require an outgroup to call substitutions, Tajima’s *D* ignores information on changes in the density of polymorphism nearby a selected site. Despite the frequent application of these two statistics, neither is specifically designed for long-term balancing selection, and both have been shown to have limited power under such scenarios (DeGiorgio et al., 2014; Siewert and Voight, 2017; Bitarello et al., 2018).

There has been a recent surge in the development of methods for specifically identifying signatures of long-term balancing selection (DeGiorgio et al., 2014; Gao et al., 2015; Siewert and Voight, 2017; Bitarello et al., 2018). Based on the Kaplan-Darden-Hudson model (Kaplan et al., 1988; Hudson and Kaplan, 1988), DeGiorgio et al. (2014) presented a mechanism to compute probabilities of polymorphism and substitution under long-term balancing selection, and developed the likelihood ratio test statistics *T*_1_ and *T*_2_. The latter statistic uses both polymorphism density and allele frequency information, and exhibits higher power than a number of methods (DeGiorgio et al., 2014; Siewert and Voight, 2017). Complementary to model-based methods, whose high power partly relies on sophisticated data, novel summary statistics have also recently been developed for detecting long-term balancing selection. Notably, two new summary statistics,β (Siewert and Voight, 2017) and non-central deviation (NCD; Bitarello et al., 2018), have been proposed to capture departures of allele frequencies in a genomic window from a certain equilibrium frequency, and both have been demonstrated to outperform the HKA and Tajima’s *D* statistics. Moreover, β, which adopts a formulation similar to Tajima’s *D*, regards the equilibrium frequency as that of the polymorphic site that a window is centered on, whereas NCD takes a user-assigned value. Further, while β does not benefit much from incorporating sites of substitution (Siewert and Voight, 2017), Bitarello et al. (2018) have shown that NCD exhibits substantially higher power and outperforms when substitutions are provided in addition to polymorphisms.

However, while several key examples (Klein et al., 1998; Ségurel et al., 2012; Leffler et al., 2013; Teixeira et al., 2015) have illustrated that it is possible and potentially common for long-term balancing selection to be established prior to speciation events, few extant approaches address the issue of identifying loci under balancing selection shared by multiple species via genome-wide scans. Traditionally, polymorphisms shared across species have been used in many studies as a tell-tale sign of shared long-term balancing selection (*e.g*., Takahata et al., 1992; Klein et al., 1998; Cho et al., 2006; Ségurel et al., 2012; Leffler et al., 2013), as they are highly suggestive of its footprints (Wiuf et al., 2004). However, such co-incident (*i.e*., trans-species) polymorphisms can also result from high mutation rates or technical artifacts introduced in sequencing and mapping. Gao et al. (2015) addressed this issue by deriving, under the Kaplan-Darden-Hudson model, the length of the balanced ancestral segment, the expected number of trans-species polymorphisms, as well as the extent of linkage disequilibrium between such polymorphisms. Although powerful, this framework is difficult to extend to an arbitrary number of species, and also requires that such trans-species polymorphisms are not missing from the dataset due to sampling or filtering. To circumvent some of these hurdles, it would be useful to be able to uncover footprints of ancient trans-species balancing selection by only using between-species substitutions and within-species polymorphisms, rather than trans-species polymorphisms.

In this study, we present a number of summary- and model-based approaches for detecting ancient trans-species balancing selection that do not rely on trans-species polymorphisms. In particular, we adapted the framework of DeGiorgio et al. (2014) to construct likelihood ratio test statistics, *T*_1,trans_ and *T*_2,trans_, to detect trans-species balancing selection (see *Theory* and *Supplementary Note* 5). Moreover, we modified the HKA test so as to better accommodate genomic data of multiple species (denoted as HKA_trans_; see *Supplementary Note* 1), and extended the NCD statistic to NCD_trans_ (see *Supplementary Note* 2) for application on multi-species data. Note that although the β statistic has been demonstrated to perform well in detecting ancient balancing selection in single species, it does not benefit from the inclusion of substitutions (as shown by Siewert and Voight, 2017), which can be highly informative when analyzing population-scale genomic data of multiple species. We therefore chose to only consider single- and trans-species variants of *T*_1_, *T*_2_, HKA (summary statistic analogue to *T*_1_), and NCD (summary statistic analogue to *T*_2_). We performed extensive simulations to evaluate the performances of these methods, and addressed a variety of confounding factors such as recent population size changes, inaccurate recombination maps, elevated mutation rates, convergent balancing selection, and window size used to compute each statistic. Next, we applied the model-based *T*_2,trans_ statistic to whole-genome human and chimpanzee data to gain insights on ancient balancing selection affecting these lineages. Further, so that these multi-species statistics can be readily applied by the scientific community, we implemented the model-based and summary statistic approaches into new software packages *MULLET* (MULti-species LikElihood Tests) and *MuteBaSS* (MUlTi-spEcies BAlancing Selection Summaries), respectively.

## Theory

Given a sample of *n* lineages, the equilibrium frequency *x* of a balanced polymorphism, and population-scaled recombination rate *ρ* = 2*Nr* between a focal neutral site and a putative selected site, DeGiorgio et al. (2014) demonstrated how the expected tree height *H*(*n, x, ρ*) and expected tree length *L*(*n, x, ρ*) of a neutral genealogy linked to a site under strong balancing selection can be efficiently calculated. These quantities can be utilized to identify genomic segments undergoing ancient balancing selection by using polymorphism and divergence data in a pair of species.

### Detecting trans-species balancing selection on two species

Consider polymorphism data from a pair of species, 1 and 2, in which we have obtained sites that are polymorphic only in species 1, polymorphic only in species 2, and substitutions (fixed differences) between the species. Suppose that at a site in the genome, the number of lineages sampled from species 1 is *n*_1_ and the number from species 2 is *n*_2_. Denote the collection of sample sizes for the two species at a locus by n = [*n*_1_, *n*_2_]. Further, suppose that by using genome-wide data, the estimated coalescence time between the two species is 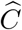. Assuming a site is *ρ* recombination units from a site with alleles maintained at frequencies *x* and 1 − *x* through strong balancing selection, analogous to the computations of DeGiorgio et al. (2014), the probability (Figure 1) of observing a polymorphic site only in species *k*, *k* = 1 or 2, is

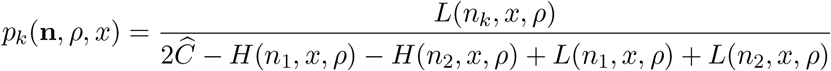

and the probability of observing a substitution between the species is

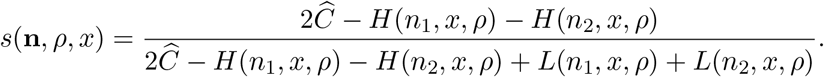

Note that *s*(n, *ρ, x*) = 1 *− p*_1_(n, *ρ, x*) *− p*_2_(n, *ρ, x*), and that our model assumes that species 1 and 2 are reciprocally monophyletic.

**Figure 1:**
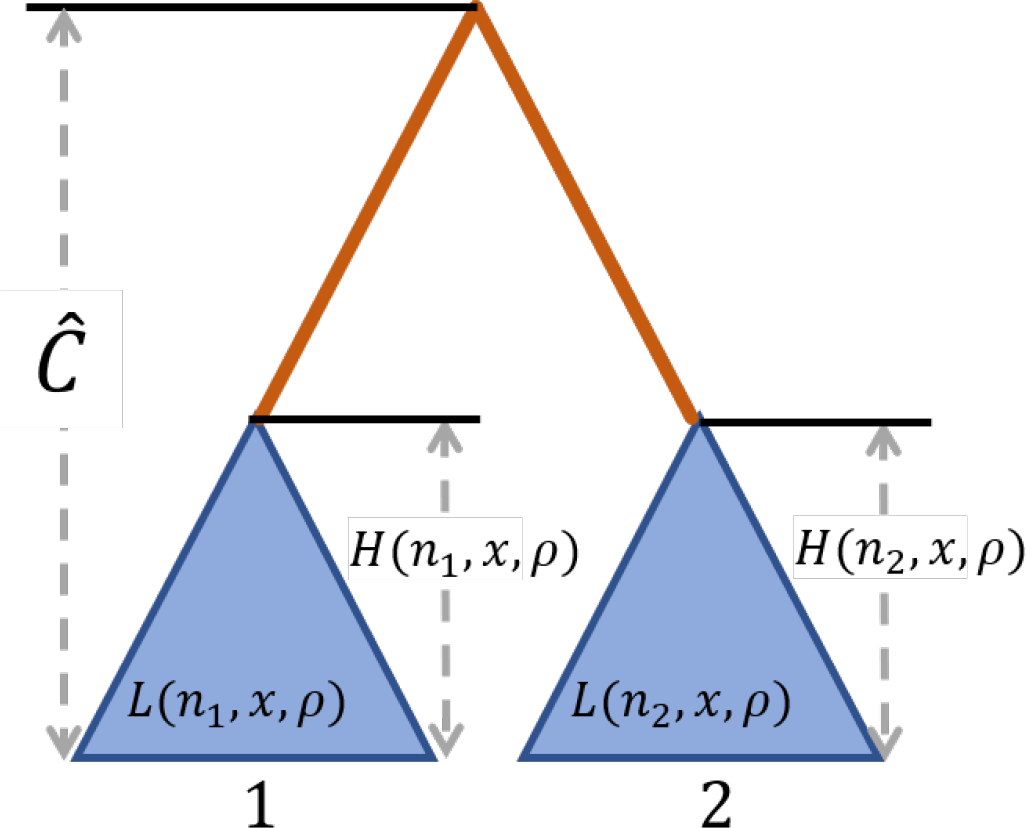
Schematic of the procedure for computing probabilities of polymorphism and substitution for a pair of species, under a model of long-term balancing selection. Blue triangles represent the subtrees of the neutral genealogy comprised of all sampled lineages for each species, where within-species polymorphic sites are generated. The orange line, which has length 2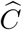 − *H*(*n*_1_, *x, ρ*) *− H*(*n*_2_, *x, ρ*), represents the length of the branch separating the two species, where substitutions are generated. 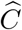 denotes the estimated expected coalescence time between species 1 and 2. *H*(*n, x, ρ*) is the expected height of the subtree for a site with *n* alleles observed that is *ρ* population-scaled recombination units from a site undergoing long-term balancing selection and maintaining alleles at frequencies *x* and 1 *− x*. *L*(*n, x, ρ*) is the expected length of this subtree.

### A composite likelihood ratio test

We can use these probabilities of polymorphism and substitution to create a likelihood ratio test for detecting trans-species balancing selection. Consider a genomic window containing *I* informative sites, where an informative site is a polymorphism only in species 1, a polymorphism only in species 2, or a substitution. Let the sample sizes in species 1 and 2 at site *i*, *i* = 1, 2, …, *I*, be *n_i_*_1_ and *n_i_*_2_, respectively. Further, suppose the derived allele counts in species 1 and 2 at site *i* are *a_i_*_1_ and *a_i_*_2_, respectively. Moreover, assume site *i* is *ρ_i_* recombination units from a site that we believe to be undergoing strong balancing selection, maintaining a pair of alleles at frequencies *x* and 1 − *x* in the population. We call this site under selection our test site. Assuming that n_*i*_ = [*n_i_*_1_, *n_i_*_2_] and a_*i*_ = [*a_i_*_1_, *a_i_*_2_], we define the vectors N = [n_1_, n_2_, …, n_*I*_], A = [a_1_, a_2_, …, a_*I*_], and *ρ* = [*ρ*_1_, *ρ*_2_, …, *ρ_I_*] for convenience. Under the alternative hypothesis of balancing selection, the composite likelihood that the test site is undergoing strong long-term balancing selection is

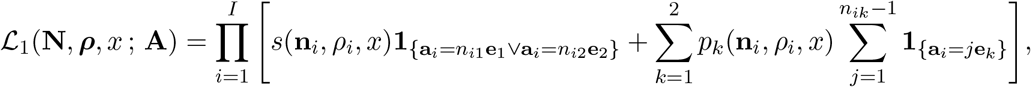

where e_*k*_ is the two-dimensional standard basis vector in which all elements are 0 except for the *k*th element, which is 1. This likelihood function is maximized at 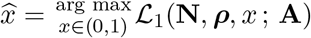.

If we condition only on informative sites, then denote the proportion of such sites across the genome that are polymorphic only in species *k* with derived allele count *a*, 0 *< a < n_k_*, by 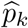(n, *a*), and the proportion of such sites that are substitutions between the species by 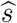(n). From these definitions, it follows that the proportion of informative sites that are polymorphic only in species *k* is 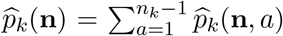. Under the null hypothesis of neutrality, the composite likelihood that the test site is evolving neutrally is

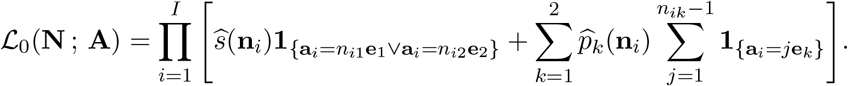

Based on the likelihoods under the null hypothesis of neutrality and the alternative hypothesis of trans-species balancing selection, we can compute a log composite likelihood ratio test statistic that the test site is undergoing strong long-term trans-species balancing selection as

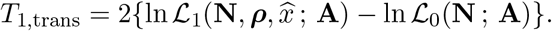

### A composite likelihood ratio test using the allele frequency distribution

DeGiorgio et al. (2014) demonstrated that such composite likelihood approaches can exhibit greater power if information on the frequency spectrum is used in addition to information on the proportions of polymorphisms and substitutions in a genomic region. For a sample of *n* alleles, conditioning on a mutation leading to a polymorphic site and assuming a locus undergoing strong balancing selection maintaining a pair of alleles at frequencies *x* and 1 − *x*, the probability under the Kaplan-Darden-Hudson model (Kaplan et al., 1988; Hudson and Kaplan, 1988) of observing a derived allele with count *a*, 0 *< a < n*, at a neutral locus *ρ* recombination units away from the selected site is *f*(*n, a, ρ, x*). Calculating *f*(*n, a, ρ, x*) analytically is non-trivial, and we instead utilize a set of frequency spectra simulated under the Kaplan-Darden-Hudson model as in DeGiorgio et al. (2014). Using the knowledge of this distribution of allele frequencies, we can modify the likelihood under the alternative hypothesis of trans-species balancing selection by including information on the frequency spectrum as

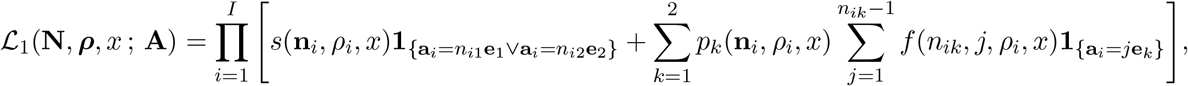

which is maximized at 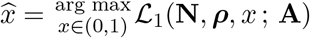

Analogous to the case without the distribution of allele frequencies, under the null hypothesis of neutrality, the composite likelihood that the test site is evolving neutrally is

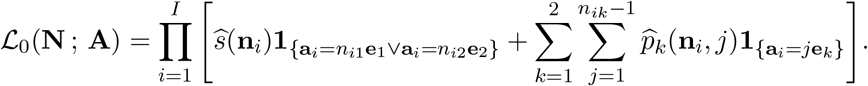

Based on the likelihoods under the null hypothesis of neutrality and the alternative hypothesis of trans-species balancing selection, we can compute a log composite likelihood ratio test statistic that the test site is undergoing strong long-term trans-species balancing selection as

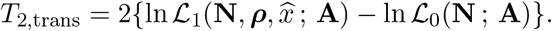

## Results

To evaluate the performances of both extant and novel methods for detecting long-term balancing selection, we chose to consider statistics that operate on the same set of informative sites, which are comprised of within-species polymorphisms and cross-species substitutions. Specifically, we considered single- and trans-species variants of *T*_1_ and its summary statistic analogue HKA (see *Supplementary Note* 1), as well as *T*_2_ and its summary statistic analogue NCD (see *Supplementary Note* 2). Considering there are eight different statistics to compare, hereafter we will refer to those designed for two or more species as “trans-species methods”, and the original variants as “single-species methods”. For empirical application, we applied the trans-species variant of the *T*_2_ statistic (*T*_2,trans_) to whole genome data from humans (The 1000 Genomes Project Consortium, 2015) and chimpanzees (Auton et al., 2012) to examine ancient balancing selection affecting both great ape species.

### Evaluating method performance on simulated data

We employed the forward-time simulator SLiM (Messer, 2013) to generate sequences of length 50 kilobases (kb) evolving along a three-species phylogeny (Figure 2; see *Materials and Methods*), under diverse selection scenarios with varying selection strength (*s*), dominance parameter (*h*), and selected allele age, as well as confounding factors such as population size changes, skewed recombination rates, and elevated mutation rates. Because footprints of ancient balancing selection are typically narrow (Hudson and Kaplan, 1988; Charlesworth, 2006; Leffler et al., 2013; Siewert and Voight, 2017), and considering that summary statistics for detecting such footprints often reach optimal performances when utilizing data from neighboring regions of similar length, we adopted a window size of one kb for single- and trans-species variants of HKA and NCD when applying them on simulated data to optimize their performances (see Figure S1 for their performances using window sizes of 0.5, 1, 1.5, 2, 2.5 and 3kb. In particular, the footprints of long-term balancing is in theory 1/(4*N_e_r*), which equals 2.5 kb when assuming a recombination rate of *r* = 10^−8^ per site per generation and an effective population size of *N_e_* = 10^4^. To ensure one kb is indeed a better choice of window size than 2.5 kb, we tested and compared their power for detecting balance selection of varying age under three distinct demographic models (Figures 2 and S2, for methods using one-kb and 2.5-kb windows, respectively). To match the amount of data available at each step (*e.g*., DeGiorgio et al., 2014; Siewert and Voight, 2017), we performed scans with the *T*_1_ and *T*_2_ variants with 10 informative sites on either side of the test site (see *Supplementary Note* 3).

**Figure 2:**
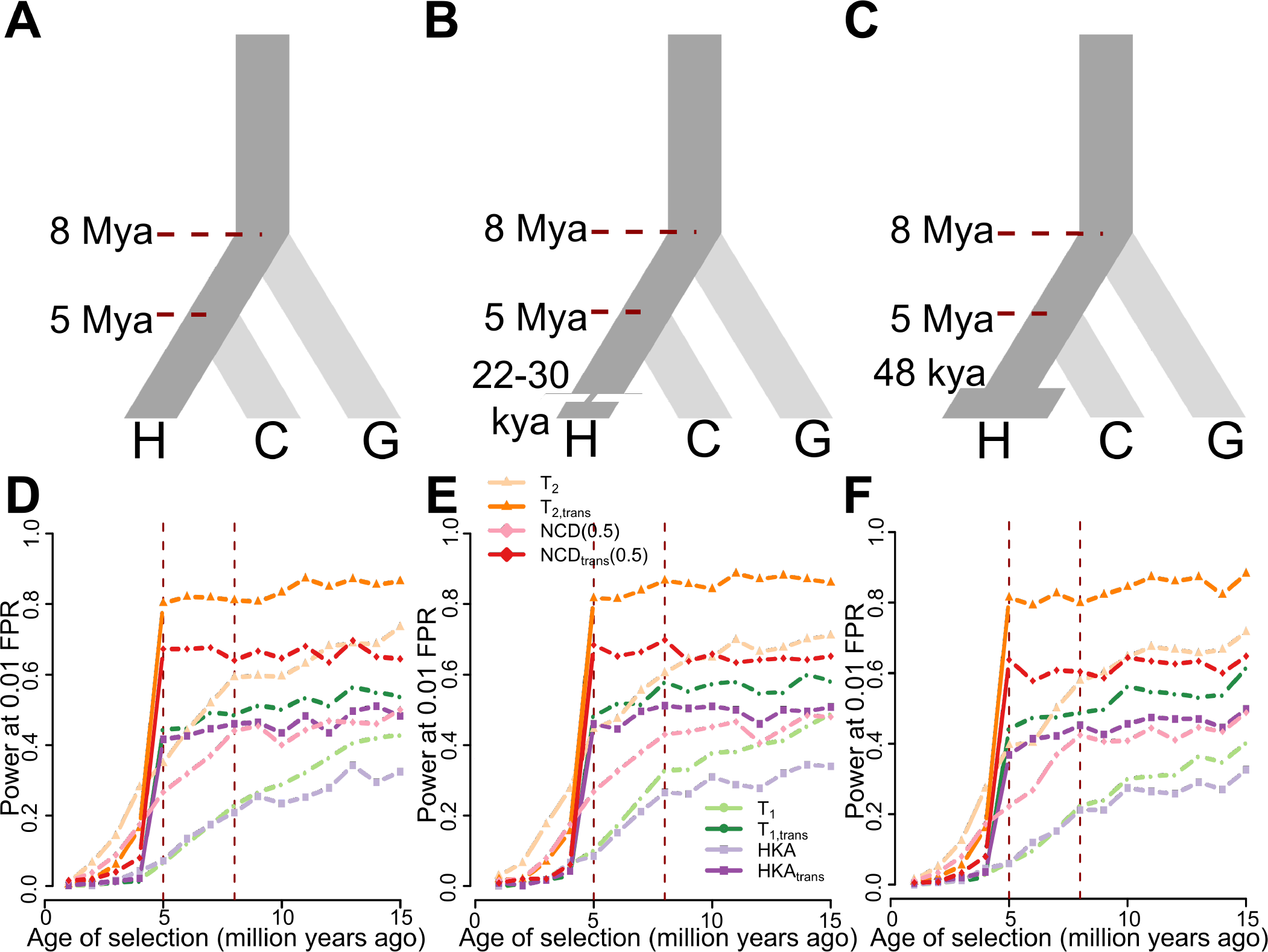
Performances of single- and trans-species variants of HKA, NCD(0.5), *T*_1_, and *T*_2_. (A-C) Schematic of demographic models relating three species, representing human (species H), chimpanzee (species C), and gorilla (species G), adopted in simulations. (A) All three species maintain constant population size of *N* = 10^4^ diploid individuals, with species H diverging from species C five million years ago (Mya), and the common ancestor of species H and C diverging from species G eight Mya. (B) Species H went through a 400-generation population bottleneck with size *N_b_* = 550 diploids 22 to 30 thousand years ago (kya). (C) Species H doubled its population size to *N_e_* = 2 × 10^4^ diploids 48 kya. Simulations assumed a generation time of 20 years across the entire phylogeny. (D-F) Powers at a 1% false positive rate (FPR) of single- and trans-species variants of HKA, NCD(0.5), *T*_1_, and *T*_2_ to detect balancing selection (*s* = 0.01 with *h* = 100) of varying age under (D) constant population size, (E) recent strong population bottleneck, and (F) recent population expansion scenarios. Red vertical dashed lines represent the times at which species H and C split, and at which species G split, respectively.

### High power of trans-species methods for detecting shared balancing selection

To assess the performance of each method in detecting balancing selection of varying age, we first modeled heterozygote advantage with selective coefficient *s* = 0.01 and dominance coefficient *h* = 100 at a genomic position, with the selected allele introduced at varying time points along the branch ancestral to species H under a demographic history of constant population size (Figure 2A). We then evaluated the powers at a 1% false positive rate (FPR) for each method. All single-species methods gain power with increasing age at which the selected allele was introduced (Figure 2D). For example, the power of *T*_2_ for detecting balancing selection has increased from almost zero to over 0.7 when the age of selection increased from one to 15 million years. Meanwhile, all trans-species methods have minimal power to detect selection established after species H and C diverge, and exhibit a surge of power for balancing selection predating the species split (Figure 2D). This jump in power is both expected and sensible because trans-species methods take polymorphism data from both species into consideration, and the adaptive changes in only one of the two should not be conspicuous enough for trans-species methods. Additionally, when balancing selection was established prior to species divergence, all trans-species methods show higher power than methods designed for single species, likely due to the greater amount of information used in their inferences.

For methods designed to operate on the same number of species, their relative performances follow similar orders. Both only utilizing polymorphism density data, the HKA and *T*_1_ variants show comparable powers (Figure 2D). Integrating allele frequency information in addition to polymorphism density data, both *T*_2_ and NCD variants outperform the *T*_1_ and HKA variants. Moreover, the *T*_2_ variants exhibit substantially higher power than the NCD variants, which is sensible given that *T* statistics are based on an explicit model and take distances between informative sites and the test site into consideration. The superior performance of *T*_2_ variants over other approaches remains consistent across varying selection strengths *s* and dominance coefficients *h* (Figures S3A, D, and G), and both display decreased power as selective advantage of heterozygotes (*i.e*., composite parameter *hs*) decreases. *T*_1_ and HKA still have similar powers, and both perform better when *hs* decreases. Meanwhile, the close margin between the HKA and *T*_1_ variants is probably because HKA and HKA_trans_ were given optimal window sizes, whereas *T*_1_ and *T*_1,trans_ may be able to effectively use information outside of the window size it was given. When the window size increases (Figure S4), *T*_1_ variants exhibit higher power than HKA variants.

Recent collapses or expansions in population size can also affect the allele frequency spectrum, as well as polymorphism density, and therefore potentially confound inferences of balancing selection. To test the robustness of each method to recent population size changes, we simulated models with a recent bottleneck (Figure 2B) or expansion (Figure 2C), using parameters inferred by Lohmueller et al. (2009) (see *Materials and Methods*). We first tested the robustness of each statistic to falsely attributing effects of population bottlenecks or expansions as footprints of balancing selection (Figure S5). These results illustrate that all methods are robust to neutral regions that are affected by strong recent bottlenecks (Figure S5A) and by recent expansions (Figure S5B). When species H underwent a recent bottleneck (Figure 2E) or expansion (Figure 2F), we observed that all trans-species methods still maintain high power to detect balancing selection whose onset predated the species divergence, outperforming their single-species counterparts. Moreover, their specificity for trans-species balancing selection also remained unaltered, and these properties also persist across different selection parameters (Figure S3).

### Distinguishing ancestral and convergent balancing selection

The trans-species statistics we developed for shared balancing selection evaluate whether there exists an increased density of polymorphic sites, an enrichment of middle-frequency alleles at polymorphic sites, or both, and whether this pattern is shared across the species examined. Because they do not directly address whether balancing selection on a candidate site predates speciation as would that of Gao et al. (2015), the statistics may be sensitive to shared, non-ancestral (*i.e*., convergent) balancing selection.

To test whether our trans-species methods can distinguish ancestral from convergent balancing selection, we simulated three scenarios in which a mutation under selection (*s* = 0.01 with *h* = 100) was introduced in one species (Figures S6A and D), two selected mutations were introduced independently in both sister species at the same site (Figures S6B and E), and two selected mutations were introduced independently in both sister species but at different sites that are 10 kb apart (Figures S6C and F). Unsurprisingly, all trans-species statistics exhibit substantially higher power than single-species statistics when the balanced alleles independently arose in both species at the same site (Figures S6B and E), compared to when only one species was affected (Figures S6A and D). This result suggests that convergent balancing selection can indeed leave similar footprints as ancestral balancing selection, provided it is old enough (four million years, or 2 × 10^5^ generations in our simulations), and acts on the same site in both species.

Nonetheless, when balancing selection in two species occurred instead at two nearby sites, the trans-species statistics (Figures S6C and F) show only moderate increases in power to falsely identify this convergent process as ancestral, and perform no better than the single-species variants. This robustness is sensible because long-term balancing selection leaves a small footprint in the genome (Hudson et al., 1987; Charlesworth, 2006; Siewert and Voight, 2017; Bitarello et al., 2018), such that the footprints around sites that are relatively close in the genome would still be unlikely to overlap and confound our trans-species methods. These results suggest that balancing selection acting on distinct sites has limited misleading effects for trans-species methods, provided the distance between these sites is larger than the expected long-term balancing selection footprint. Further, in situations where the distance between selected sites is smaller than the selection footprint, the biological mechanisms leading to their maintenance of polymorphisms are likely similar.

### Effect of error in recombination rate estimation

In addition to the improved power of trans-species methods to detect ancient balancing selection, we have demonstrated the superior specificity and robustness of the model-based *T*_2_ statistics, especially *T*_2,trans_. Nonetheless, other non-adaptive events, such as skewed recombination rates or inaccurate recombination maps, may potentially interfere with the detection of long-term balancing selection, and perhaps have a more deleterious impact on the model-based *T* statistics that rely on estimates of recombination rates. To examine method robustness to skewed recombination rates, we generated 50 kb long sequences under models of unevenly-distributed recombination rates fluctuating every one kb along the sequence, and we considered fluctuations of two different orders of magnitude. Specifically, assuming a recombination rate of *r* = 10^−8^ per site per generation under our earlier scenarios of a uniform recombination map, we set recombination rate to alternate along the sequence from 10*r* to *r*/10 (10^2^-fold change between adjacent regions; Figures S7A and C) or from 100*r* to *r*/100 (10^4^-fold change between adjacent regions; Figures S7B and D). With the correctly informed coalescence time, polymorphism-to-substitution ratio, and derived allele frequency spectra, we applied all methods on the simulated sequences, and let *T* statistics assume a uniform and constant recombination rate of *ρ* = 2*N_e_r*, as they do in all other simulation scenarios. Providing *T* statistics with such an erroneous recombination map permits us to evaluate the robustness of these statistics when the recombination rates are grossly mis-specified.

All methods are robust to falsely identifying neutrally-evolving regions with massive fluctuations in recombination rate as balancing selection (Figures S7A and B). Furthermore, for sequences with recombination rate fluctuating by 10^4^-fold (Figure S7B), the proportion of false signals for each method further decreases, most outstandingly for *T*_2_ and *T*_2,trans_. On the other hand, when an allele under balancing selection (*s* = 0.01 with *h* = 100) was introduced in the ancestral population 15 million years ago, all methods show increased power when recombination rate fluctuates by 10^2^-fold (Figures S7C and E). When the rate fluctuates by 10^4^-fold (Figure S7D), all single-species methods show compromised power compared with those under 10^2^-fold change (Figures S7D and E). However, single- and trans-species variants of both polymorphism density-based methods, *T*_1_ and HKA, exhibit improved power under skewed recombination maps compared to those under a uniform map, whereas variants of methods that incorporate information on allele frequencies, *T*_2_ and NCD, do not always gain power (Figure S7E). Despite that *T*_2,trans_ and NCD_trans_ both have marginally higher power under skewed recombination maps, the increase in power exhibited by *T*_1,trans_ and HKA_trans_ was considerably greater. This discrepancy may be due to irregular recombination shifting the spatial distribution of polymorphic sites around a selected site, while exerting little influence on allele frequencies at these polymorphic sites.

### Robustness to high mutation rate

Another non-adaptive phenomenon that may confound inferences of ancient balancing selection is the increase in mutation rate at a genomic region. To test the robustness of each method against elevated mutation rate, we simulated sequences neutrally evolving along the demographic history in Figure 2A, and mutating at rates that are five-, 10-, or 20-fold higher than the original rate of *µ* = 2.5 × 10^−8^ per site per generation. In addition to mutational hotspots shared across all species, we also considered the scenarios in which the hotspots are species-specific. To examine the robustness of each method to species-specific elevation of mutation rates, we simulated a set of scenarios in which sequences in species H (as labeled in Figure 2A) started to mutate at rate 5*µ*, 10*µ*, or 20*µ* after splitting from species C (see *Materials and Methods*). We then applied both single- and trans-species methods on these sequences with the same window sizes used in previous power analyses so as to mimic the effect of unexpected mutation hotspots. Note that for *T* statistics, we provided estimates of coalescence time, proportions of polymorphisms and substitutions, and site frequency spectra based on neutral data generated under the original mutation rate *µ*. Similarly, we provided the HKA statistics with proportions of polymorphisms and substitutions expected from neutral data simulated with the original mutation rate *µ*.

When mutation rates were elevated across all sequences along the phylogeny, most statistics exhibited elevated proportions of false signals (Figure S8), with higher mutation rates leading to greater numbers of false signals. One exception is NCD, whose proportion of false signals at a 1% FPR remains substantially smaller than 0.01 for all scenarios with elevated mutation rates (Figure S8). This robustness of NCD results from the increased number of informative sites incorporated in each window, which leads to smaller variance of NCD scores. Among the statistics that exhibited increased mis-identification rates, *T* statistics (especially *T*_trans_) reported lower proportions of false signals than others. Meanwhile, both single- and trans-species HKA statistics exhibit considerable proportions of false signals at a 1% FPR (Figures S8D-F), even reaching 0.7 when the mutation rate elevated 20-fold (Figure S8F), which is almost twice the mis-identification rates of *T*_1_ and *T*_2_. Nonetheless, this inflation is sensible for HKA because its chi-square formulation (*Supplementary Note* 1) accounts for the number of informative sites within the window (*i.e*., number of observations), and would yield larger values when more sites are considered. Further, because HKA was not specifically designed for detecting balancing selection, but rather to identify departures from expected numbers of polymorphisms and substitutions, it is natural that it should uncover regions with higher mutation rates.

When instead the mutation rate increased only in species H, we found that all single-species methods mis-identified substantial proportions of such sequences as evolving under long-term balancing selection (Figure S9). In particular, they showed considerably inflated mis-identification rates compared with scenarios of mutation rates elevated uniformly across all species (Figure S8), among which single-species variants of *T*_1_ and HKA even reported over 90% false signals at a 1% FPR across all three elevated mutation rates considered. Meanwhile, although trans-species methods (except for NCD_trans_) displayed considerable increases in proportions of false signals under species-specific accelerated mutation (Figure S9) compared with those under uniformly elevated mutation rates (Figure S8), this inflation in false signals was substantially less dramatic than for variants of single-species statistics. This higher robustness of trans-species relative to single-species methods is consistent with their behaviors under uniform mutation rate increases (Figure S8).

Despite the large performance margin for each statistic between scenarios of shared and species-specific mutational hotspots, it is sensible that the latter would be more misleading, especially for *T*_1_ and HKA. When the mutational process is only partially accelerated along the phylogenic tree relating multiple species, the accumulation of mutations along certain lineages can lead to elevated polymorphism-to-substitution ratios. Such distortions can be observed in our simulated sequences in species H (Figure S10), which explains the drastic inflation of the false signals identified by *T*_1_ and HKA (Figure S9). Meanwhile, as expected, the allele frequency spectra are unaffected (Figures S10), thereby providing advantage to methods, such as *T*_2_ and NCD, that also employ information on allele frequencies. When data from both species are considered, however, the distortion in polymorphism density was ameliorated (Figure S11), but the proportion of polymorphisms segregating in species H drastically increased as the species-specific mutation rate increased. Considering that the overall proportions of substitutions (Figure S11) as well as allele frequency spectra remain relatively constant, this distortion in proportions of sites being polymorphic in species H makes it reasonable that trans-species methods identify more false signals than they would under uniformly high mutation rate, but are still much less affected by the species-specific mutational hotspot than were the single-species statistics. We further explore the reasons for the inflation of *T* statistic mis-identification rates under scenarios of increased mutation rate in the *Discussion* section.

### Effect of window size

For all extant methods to detect signatures of long-term balancing selection, the length of the genomic region considered (hereafter referred to as “window size”) when computing their scores could substantially impact their powers. In our study, we applied all summary statistics on windows with a fixed number of nucleotides, and *T* statistics with a fixed number of informative sites surrounding each test site. To best match the amount of data provided to *T* statistics, we set the number of informative sites flanking the test site on either side as *I*, such that 2*I* +1 is closest to the expected number of sites included in a one kb region under neutrality given the simulation parameters (see *Supplementary Note* 3). Because long-term balancing selection typically leaves behind narrow genomic footprints, *i.e*., of length approximately 1/(4*N_e_r*), where *r* is the per-site per-generation recombination rate (Hudson and Kaplan, 1988), extant methods often reach optimal power when the window size is of the same magnitude of this size, which in humans is approximately 2.5 kb, given an effective size of *N* = 10^4^ and *r* = 10^−8^. To choose the optimal window size for our analyses on simulated data, we examined the relative performances of summary statistics under 0.5, 1, 1.5, 2, 2.5, and 3 kb windows (Figure S1), and accordingly adopted the size of one kb for other simulated data. However, although their powers can be optimized by adopting window sizes close to the narrow footprint generated by long-term balancing selection, with data derived from such a limited genomic region, estimation of these statistics can be noisy and potentially misleading. In empirical applications, it may therefore be preferable to incorporate information from a wider genomic region for the estimation of these statistics to reduce stochasticity. Hence, it is important to examine how window size affects method performance.

To this end, we applied all single- and trans-species methods considered in this study to simulated data, and varied window sizes under which the statistics were calculated (see *Materials and Methods*), and compared their powers. To ensure that all methods operated on the same data, we applied the summary statistics with windows containing a fixed number of informative sites, which is how the *T* statistics are computed. We tested windows with *I* = 5, 10, 30, 50, 100, 150, 200, 250, 300, and 350 informative sites on each side of the test site (*i.e*., the site on which windows are centered). Because we are interested in ancient trans-species balancing selection, we chose to examine the scenario in which the selected allele (*s* = 0.01 with *h* = 100) arose 15 million years (assuming a generation time of 20 years) prior to sampling.

As predicted, powers of all methods drastically decrease as window size increases (Figure S12A). While powers of all other methods eventually decrease toward zero for large windows, *T*_2_ and *T*_2,trans_ still maintain considerably higher power than other methods, with powers reaching approximately 0.2 when the number of informative sites considered is larger than 200 sites on either side of the test site (*i.e*., 401 informative sites covered by each window). This contrast can be better illustrated in Figure S12B, where *T*_2_ and *T*_2,trans_ are the only two statistics that, as the window size increases, experience decay in power substantially slower than all other statistics, and stabilize at non-zero values. The relative robustness of *T*_2_ and *T*_2,trans_ is sensible because they incorporate distances from the test site of each informative site, reducing the influence of sites too far from the tested site. However, although both *T*_1_ and *T*_2_ account for the spatial distribution of informative sites, the powers of *T*_1_ and *T*_1,trans_ decrease more drastically than of *T*_2_ and *T*_2,trans_, albeit not as quickly as its summary-statistic counterpart HKA (Figure S12B). Additionally, the powers of HKA variants decay much faster than those of NCD variants as window size increases. In general, statistics based only on polymorphism density, such as single- and trans-species variants of *T*_1_ and HKA, are more vulnerable to large window sizes when compared to those accounting for allele frequency, such as single- and trans-species variants of *T*_2_ and NCD. This gap in robustness between these two classes of statistics is likely due to the large emphasis that allele frequency-based statistics place on the presence of moderate-frequency alleles, thereby requiring a larger number of additional neutral informative sites for the evidence of the signal established from these intermediate-frequency alleles to attenuate.

### Applying *T*_2,trans_ on great ape genomic data

To examine long-term balancing selection affecting both human and chimpanzee lineages, we applied *T*_2,trans_ on genomic data merged from human (108 Yoruban individuals; The 1000 Genomes Project Consortium, 2015) and chimpanzee (10 western chimpanzees (*Pan troglodytes verus*); Auton et al., 2012). Variant calls from both species were mapped to human reference genome hg19 and polarized with the matching sequence from gorilla reference genome gorGor3 (Kent et al., 2002). We retained bi-allelic sites that were polymorphic within a single species or that were substitutions between the two species, and adopted stringent filters (see *Materials and Methods*) to reduce the potential influence of technical artifacts. We then performed a whole-genome scan with *T*_2,trans_ based on an inferred human recombination map (The International HapMap Consortium, 2007), and considered 100 informative sites directly upstream and directly downstream of the test site (*i.e*., 201 informative sites in total).

To infer statistical significance of each test site, we employed the coalescent simulator *ms* (Hudson, 2002) to generate 5 × 10^7^ independent replicates of 25 kb long sequences, evolving neutrally along the inferred demographic histories of the Yoruban, chimpanzee, and gorilla lineages (Auton et al., 2012; Prado-Martinez et al., 2013; Terhorst et al., 2017, see *Materials and Methods*), with sample sizes matching the empirical data. We applied *T*_2,trans_ on the first 201 informative sites of each sequence, using the 101st site as the test site, providing a single *T*_2,trans_ score for each replicate (see *Materials and Methods*). We assigned *p*-values to each test site in the empirical scan based on the distribution of *T*_2,trans_ scores from the neutral replicates (Figure S13). To correct for multiple testing, a test site reaches genome-wide significance if its *p*-value is smaller than 0.05/10^6^ = 5 × 10^−8^, which uses the common assumption that there are approximately 10^6^ independent sites in the human genome (Altshuler et al., 2008). Thus, based on the distribution shown in Figure S13, sites with a *T*_2,trans_ score greater than 223.709 were considered statistically significant outliers.

### Significant evidence of shared balancing selection on the MHC region, *FREM3*/*GYPE* locus, and *GRIK1*/*CLDN17* intergenic region

The major histocompatibility complex (MHC) has been repeatedly demonstrated to maintain high polymorphism levels across multiple species (Takahata et al., 1992; Leffler et al., 2013). Consistent with previous evidence, the *T*_2,trans_ statistic exhibits significantly outstanding scores in the MHC region (Figure 3). A closer examination reveals that all peaks identified in the MHC region (Figure 3) locate on the genes previously-identified to exhibit signatures of long-term balancing selection (Hedrick et al., 1991; Takahata et al., 1992; Andrés et al., 2009; Sanchez-Mazas, 2007; DeGiorgio et al., 2014; Siewert and Voight, 2017; Bitarello et al., 2018). Specifically, across the region (Figure 3) approximately four clusters of genes exhibit prominent scores, with large peaks over *HLA-A*, over *HLA-B* and *HLA-C*, over *HLA-DRB* genes, and over *HLA-DPB* genes. This pattern is consistent with the one reported by DeGiorgio et al. (2014), where these regions were extreme outliers in the scan for long-term balancing selection in human populations. Note-worthily, we observed that the most outstanding signal within this region falls on the gene *HLA-DPB1* and its pseudogene *HLA-DPB2* (Figure S14), with the former making up the beta chain of the MHC II molecules. The beta chain of MHC II is responsible for presenting extracellular immunogens, and contains polymorphisms that diversify peptide-binding specificity (Díaz et al., 2003). These results agree with the observations that polymorphisms in the MHC region are often shared among species (Takahata, 1993), and have been maintained since before the split of multiple great ape species (Takahata et al., 1992; Meyer et al., 2017).

**Figure 3:**
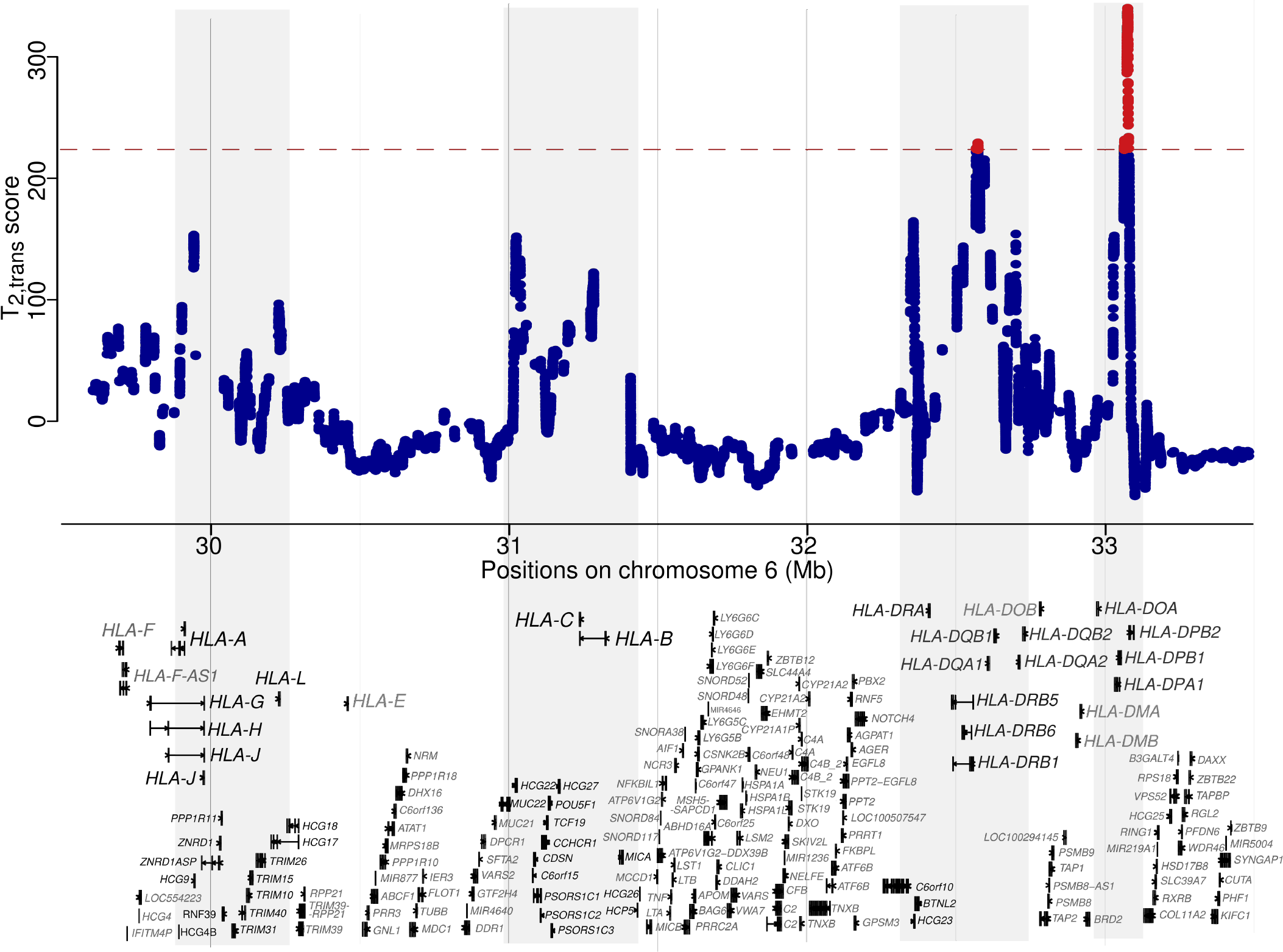
*T*_2,trans_ scores within the MHC region on chromosome 6. Gene tracks are displayed below, with only the longest transcript of each gene shown. If a gene does not have a longest transcript to cover all exons, then the minimum number of transcripts are displayed. Regions enriched with high-scoring peaks are shaded in gray, and genes falling in these areas are labeled with black fonts. The red dashed line represents the cutoff value for statistical significance, and the test sites with significant scores are shown in red.

In addition to the MHC region, we observed similarly extraordinary signals on chromosome 4, between the genes *FREM3* and *GYPE* (Figure S15). This locus was previously reported by Leffler et al. (2013) to harbor trans-species polymorphisms shared by humans and chimpanzees, and is functionally associated with malaria defense (Malaria Genomic Epidemiology Network, 2015; Leffler et al., 2017). More specifically, the polymorphisms on this locus exist in accordance with the copy number variation of *GYPB* and *GPYA*, which can result in polymorphism of a blood-group antigen that effectively defends malaria infection (Leffler et al., 2017). Interestingly, while both the MHC and *FREM3*/*GYPE* regions exhibit an enrichment of polymorphic sites (Figures S14D and S15D), we only observed a specific enrichment of minor allele frequency on the *FREM3*/*GYPE* locus, where a 400 kb-long region surrounding the peak still exhibits a distinguishable enriched frequency of approximately 0.3 (Figures S15B and C). In the MHC region, however, even after narrowing the range considered down to 200 kb around its largest peak, multiple modes can still be observed (Figures S14B and C), suggestive of complex balancing selection processes operating on this region and matching the footprints of multiple balanced loci with different equilibrium frequencies (Meyer et al., 2017).

Another significant candidate region falls on chromosome 21, between the genes *GRIK1* and *CLDN17* (Figure 4). A number of transcription factor binding sites locate on the peak region, binding the factors CTCF, RAD21, and FOS (data from Ziller et al., 2013, as the ENCODE transcription factor ChIP-seq track shown on the UCSC genome browser). Although the regulatory activity of this intergenic region still remains elusive, the genes surrounding the region have potentially intriguing functional implications. Upstream of the peak locates a kainate-selective glutamate receptor gene *GRIK1*, which has been associated with epilepsy (Sander et al., 1997) and schizophrenia (Shibata et al., 2001). Mice knocked-out of *GRIK1* would exhibit decreased pain perception (Gardiner and Costa, 2006), implying that the fine-tuning of its expression may be important for an appropriate level of perceptional acuity. On the other side of the peak, the *CLDN17* gene encodes claudin-17, which forms anion-selective channels on tight-junction barriers. It is highly expressed in kidneys and is hypothesized to be involved in chloride re-uptake (Krug et al., 2012). It is also expressed in intestine and the brain (Lonsdale et al., 2013), potentially contributing to the integrity of important barriers such as intestine and blood-brain barriers. Similarly, claudin-8, encoded by the nearby *CLDN8* gene, is also involved in chloride resorption in kidneys (Hou et al., 2010). It is an integral part of the intestine barrier (Groschwitz and Hogan, 2009), and studies have associated the gene with inflammatory bowel diseases (Zeissig et al., 2007). Moreover, claudin-8 has also been reported to be susceptible to gut bacteria endotoxin (Shrestha and McClane, 2013). To make the case more intriguing, there seems to be another high-scoring region upstream of the two claudin genes (Figure 4A), which is in the vicinity of a cluster of genes encoding hair keratin-associated proteins (Shibuya et al., 2004). As if corresponding to the two peaks, the site frequency spectra of both human and chimpanzees (Figures 4B and C) show enrichment of two different frequencies, suggesting that two distinct equilibrium frequencies may have been maintained if balancing selection were acting on this region.

**Figure 4:**
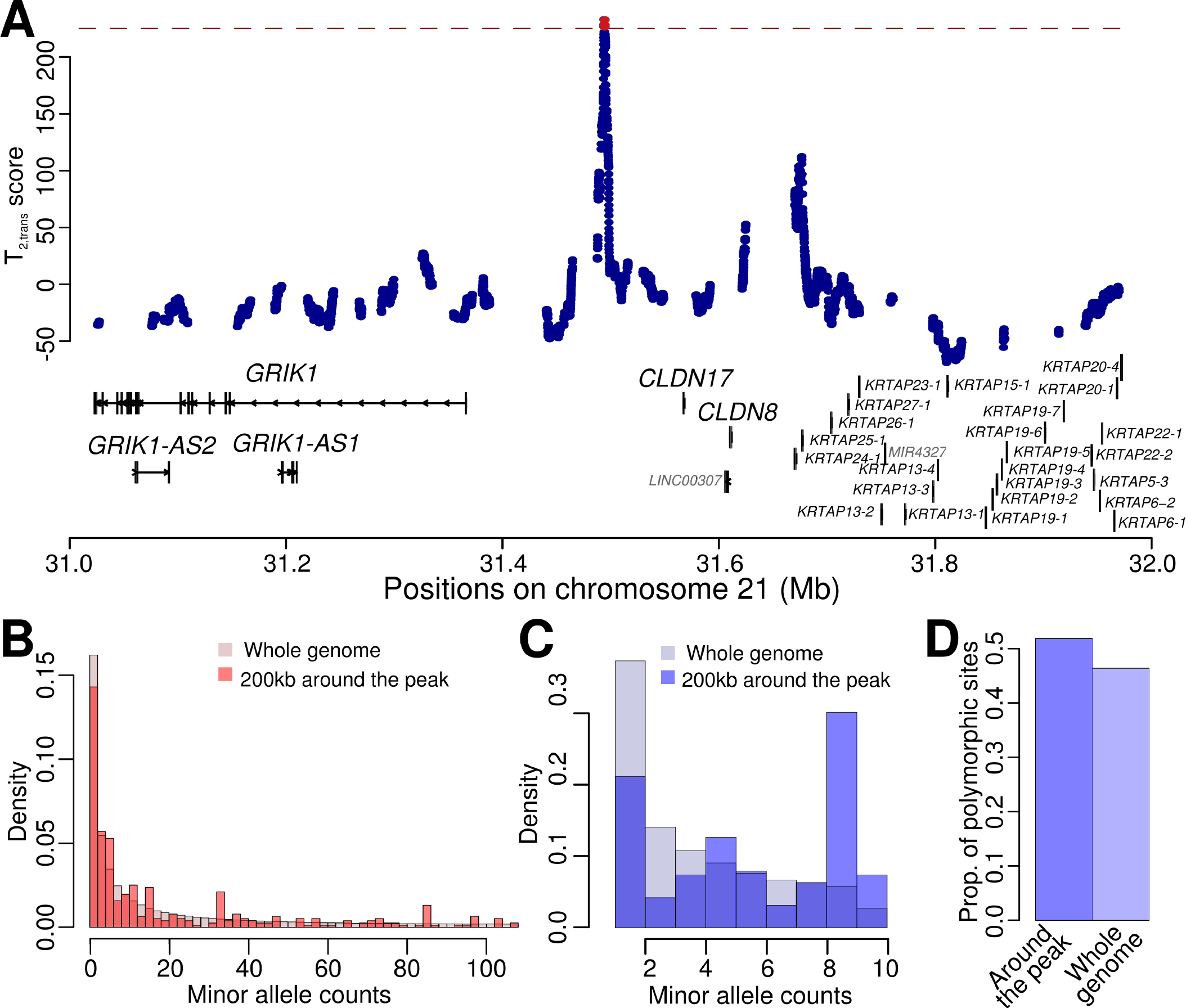
Patterns consistent with long-term balancing selection within the *GRIK1*/*CLDN17* intergenic region. (A) *T*_2,trans_ scores within the one Mb genomic region on chromosome 21 encompassing the *GRIK1*/*CLDN17* intergenic region. Gene tracks are shown on the corresponding location, with key genes labeled with larger fonts. The red dashed line represents the cutoff value for statistical significance, and the test sites with significant scores are shown in red. (B-D) Minor allele frequency spectra at polymorphic sites for human (B) and chimpanzee (C), and proportion of informative sites that are polymorphisms (D) within a 200 kb region encompassing the *GRIK1*/*CLDN17* intergenic region, compared with those of the whole genome.

### Multiple genes with outstanding evidence of trans-species balancing selection

Because we took a conservative approach to obtain *p*-values of test sites (see *Discussion*), few outstanding peaks are identified as significant genome-wide. However, some of them do show top scores that are close to the genome-wide significance cutoff, and locate on genes with potentially important functional implications, including *SLC35F1* (Figure S16) and *ABCA13* (Figure S17). Among the regions that did not meet the genome-wide significance threshold, the highest-scoring candidate is *SLC35F1*, which is located on chromosome 6 and encodes a putative ion channel. Although no functional study has been reported on this gene, multiple studies have associated it with resting heart rate levels (Pfeufer et al., 2009; Den Hoed et al., 2013), risk of atrial fibrillation (Christophersen et al., 2017), and heart attack risks (van der Ende et al., 2018). However, despite its associations with cardiac health, *SLC35F1* is mainly expressed in brain tissues, especially the cerebral cortex (Uhlén et al., 2015). Additionally, while the SNPs previously associated with cadiac functions mostly enrich under the minor peak around position 118.6 Mb of chromosome 6 (Figure S16A; eQTL data from Lonsdale et al., 2013; Ziller et al., 2013), the major peak on this gene locates around position 118.3 Mb on chromosome 6 within a cluster of DNAaseI sensitive loci (Ziller et al., 2013) in the first intron on *SLC35F1*. These results suggest a possible role of selection on regulatory function and potential pleiotropy. Another candidate gene *ABCA13*, the longest in its gene family, harbors a region with outstanding *T*_2,trans_ scores (Figure S17). This ATP-binding cassette (ABC) transporter gene is highly expressed in multipotent adult progenitor cells (MAPC; Tang et al., 2010), a rare type of multipotent stem cells that can differentiate into not only mesodermal, but also endodermal and ectodermal cells (Hof et al., 2007), and are important for wound-healing and tissue repair (Reyes and Verfaillie, 2001). In addition to bone marrow (Uhlén et al., 2015), where most blood stem cells are found, expression of *ABCA13* can also be detected in tracheae, thymus, testes, and ovaries (Prades et al., 2002; Barros et al., 2003). Moreover, the peak within this gene sits immediately upstream of the trans-membrane ABC2 domain (Figure S17), suggesting a potential selective force to diversify either the splicing or the functionality of this domain, which may be sensible given the wide variety of cell lineages that MAPC can differentiate into. Although functional studies of *ABCA13* are lacking, both its expression pattern and the location of the peak present an intriguing case for a potential target of long-term balancing selection.

### Extending extant frameworks to *K* > 2 species

So far we have demonstrated that the two-species versions of the *T*_1_, *T*_2_, HKA, and NCD statistics are specifically-tailored to detect trans-species balancing selection, display substantially higher power than single-species statistics, and can recover well-characterized cases of balancing selection shared between the human and chimpanzee lineages. Furthermore, as described in *Supplementary Notes* 1, 2, and 5, all extant frameworks can be extended to an arbitrary number of species *K*.

To test the performances of these *K*-species extensions, we simulated 50 kb long sequences over a five-taxon tree (Figure 5A), which in addition to the three-species (species H, C, and G) examined in earlier simulations, features a fourth and fifth species diverging from the others 12 and 17 million years ago, respectively, analogous to that of orangutans (denoted by species O) (Scally et al., 2012) and gibbons (denoted by species B) (Carbone et al., 2014). All other parameters remained the same as for the three-species tree with constant population sizes (see *Materials and Methods*). We introduced a selected mutation with strength *s* = 0.01 and dominance *h* = 100 in the lineage ancestral to species H at varying time points, and evaluated the performances of each method. For each statistic, we tested their extensions for application on two, three, and four species, and traced their powers as a function of time at which the selected allele arose, in addition to their single-species variant (Figure 5).

**Figure 5:**
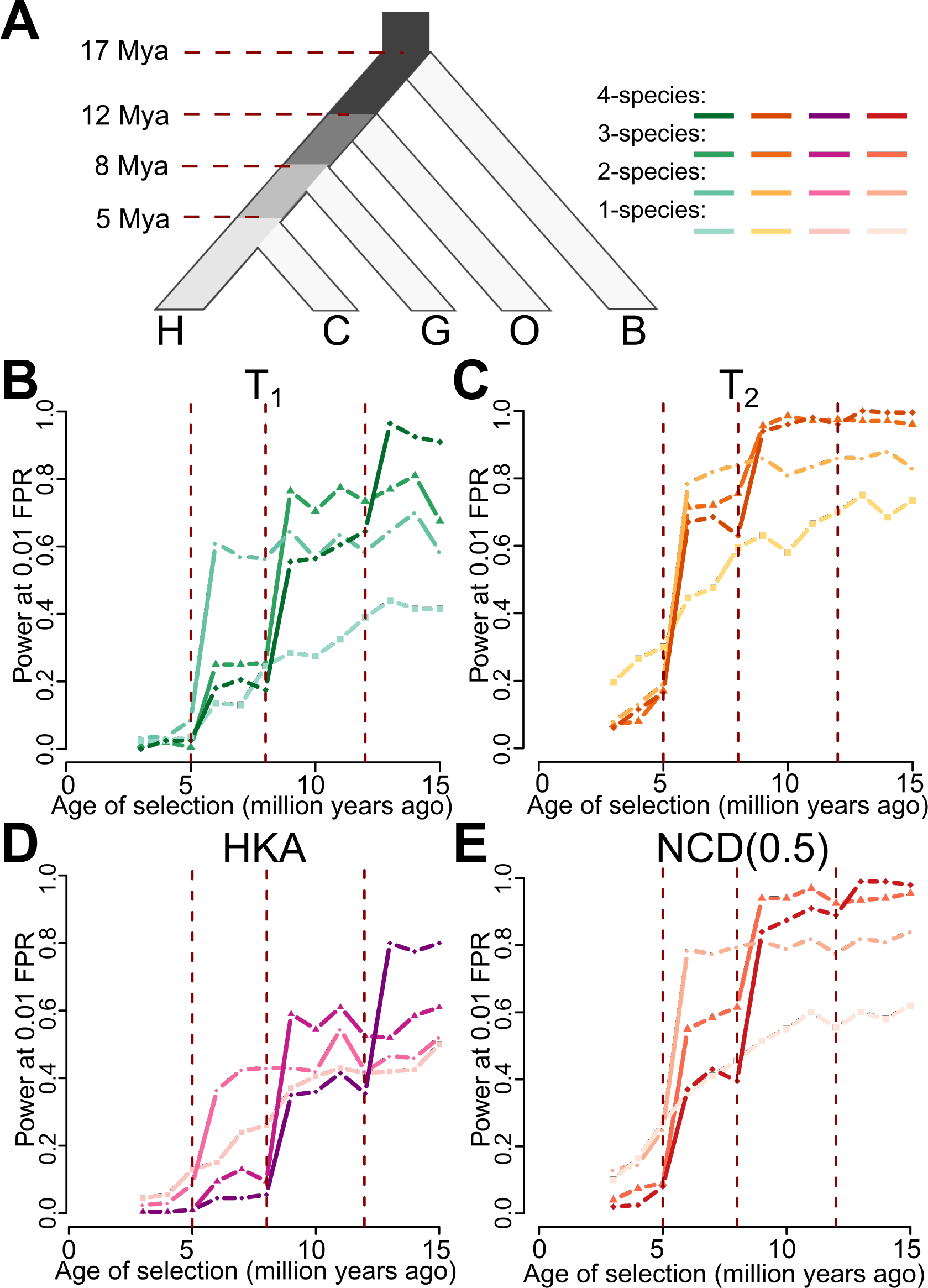
Performances of *K*-species variants of HKA, NCD(0.5), *T*_1_, and *T*_2_, with *K* = 1, 2, 3, or 4. (A) Schematic of the simulated five-species tree, relating species H, C, G, O, and B. Branches ancestral to species H are shaded based on the number of species that descend from that branch, with darker shades corresponding to larger numbers of species. (B-E) Powers at a 1% false positive rate (FPR) of one-, two-, three-, or four-species variants for (B) *T*_1_, (C) *T*_2_, (D) HKA, and (E) NCD(0.5) to detect balancing selection (*s* = 0.01 with *h* = 100) of varying age. *K*-species variants of HKA, NCD(0.5), *T*_1_, and *T*_2_ with darker shaded lines are those that consider a greater number of species *K*.

Consistent with earlier results for two-species statistics (Figure 2), all two-species statistics show substantially higher power than their single-species counterparts in uncovering balancing selection introduced prior to the split of species H and C. Similarly, all three-species statistics exhibit a surge in power once the age of balancing selection surpasses that of the divergence of species G, and all four-species statistics show an analogous increase for selection predating the divergence of species O. This relation among each *K-* species (*K* = 1, 2, 3, or 4) variant remains consistent for *T*_1_, *T*_2_, HKA, and NCD (Figures 5B, C, D, and E, respectively). Moreover, for every statistic and for each branch of interest (colored using incrementally darker shades in Figure 5) where balancing selection was introduced, the highest power can always be observed in the variant of a method that is specifically tailored for the corresponding number of species sharing the selection event. That is, for a specific method, the variant with the highest power is the one that operates on the entire set (and only this set) of species descending from a specific ancestral branch in which the selected allele arose. These results illustrate the applicability of *K*-species extensions of extant methods for detecting long-term balancing selection, and also for broadly constraining the time at which selected alleles arose at sites undergoing balancing selection.

In addition to their powers, we also examined the abilities of each method to localize the site under balancing selection (Figure 6). For each statistic, the absolute distance from their highest score to the true location decreases as the age of the balanced allele increases, consistent with the fact that older balancing selection would leave characteristically narrower genomic footprints. Within each method, each trans-species variant exhibits a steep drop in mean distance to the true site of selection once the age of selection predates the divergence of all species it examines. This improvement in accuracy accords with the sharp increase in power. As the power of each method surpases 0.8 (Figure 5), the mean distance to the true site of selection decreases to less than one kb (Figure 6). This result demonstrates that under scenarios in which methods exhibit high power to detect ancient balancing selection, their ability to isolate the true location of the selected site is considerable.

**Figure 6:**
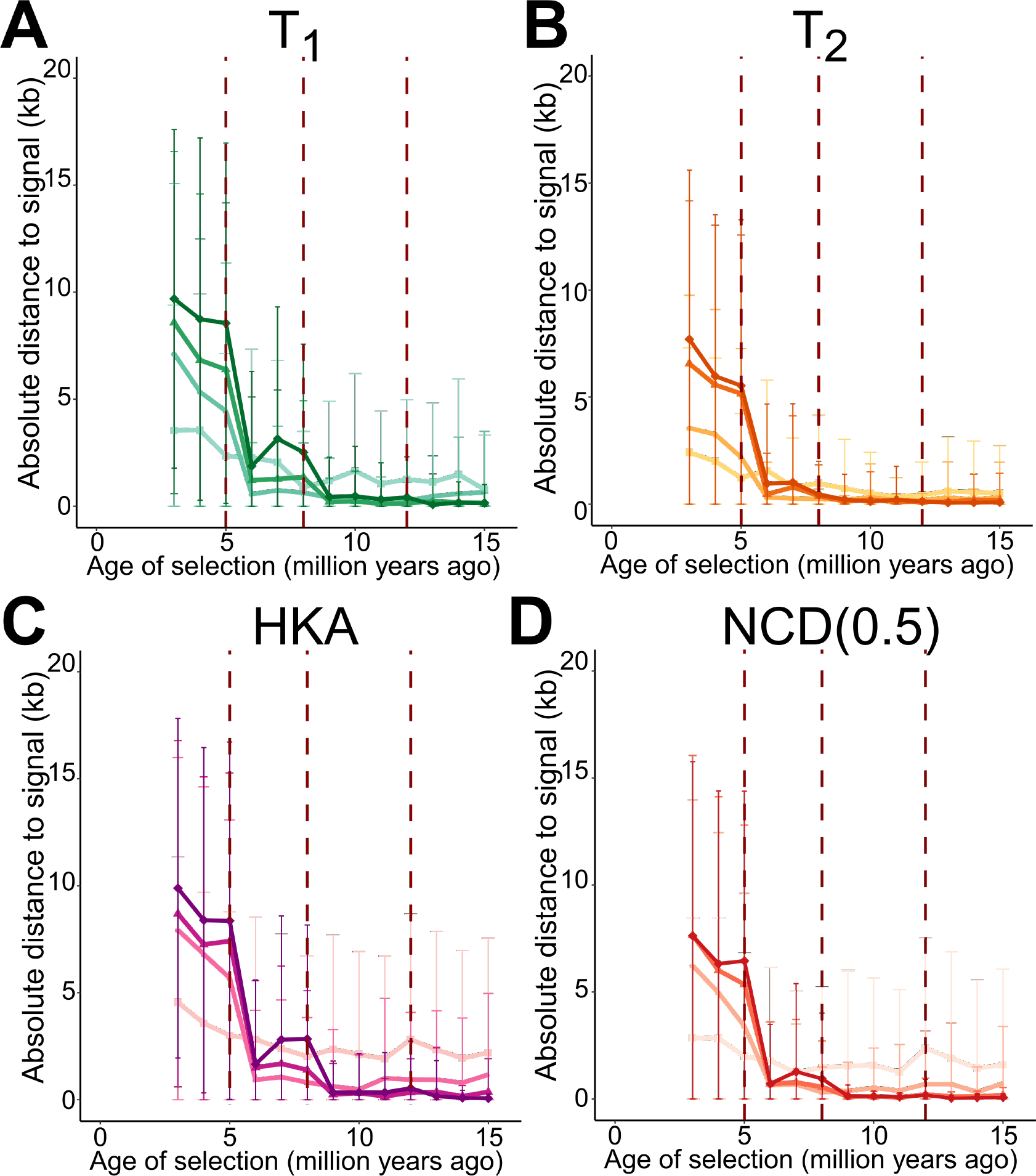
Mean absolute distances from location of signal peak to true site under balancing selection (*s* = 0.01 with *h* = 100) of varying age for *K*-species variants of (A) *T*_1_, (B) *T*_2_, (C) HKA, and (D) NCD(0.5). Error bars represent the standard deviation of all 200 replicates of the corresponding simulated scenario. Statistics are color-coded as in Figure 5.

## Discussion

In this study, we developed multi-species variants of summary-statistic and model-based approaches that employ polymorphism and substitution data from an arbitrary number of sampled species to localize sites undergoing shared balancing selection, and have comprehensively evaluated their performances through simulations. We applied the model-based *T*_2,trans_ statistic to genomic data from humans and chimpanzees, and recovered the previously-reported MHC and *FREM3*/*GYPE* regions as the most outstanding candidates. We have also characterized novel candidate regions on the cardiac health-associated *SLC35F1* and the ATP-binding cassette gene *ABCA13*, presenting intriguing cases of long-term balancing selection shared by human and chimpanzee lineages that potentially diversifies these genes’ functionality.

### Performance of trans-species methods on simulated data

In our simulation study, we demonstrated that all trans-species statistics exhibit specificity in detecting long-term balancing selection shared by multiple species. They have low power relative to their single-species variants when only a single species is undergoing balancing selection, and display high power when selection initiated prior to the divergence of all input species (Figure 2). Moreover, we tested the performances of each method under different demographic parameters (Figure 2), varying selection parameters (Figure S3), skewed recombination maps (Figure S7), and elevated mutation rates (Figures S8-S10). Our results have shown that the specificity of trans-species methods for shared balancing selection remains robust across diverse scenarios. Overall, as expected, methods using more information for the same genomic region generally attained higher power. Specifically, *T*_2_ and NCD, which consider information on allele frequencies in addition to polymorphism density, outperform *T*_1_ and HKA, which do not consider allele frequencies. Similarly, the model-based *T* statistics also outperform their corresponding summary statistic analogues (*i.e*., HKA to *T*_1_, and NCD to *T*_2_), as the model accounts for the spatial distribution of informative sites, in addition to regional polymorphism density and allele frequency distribution. We are also aware that Bitarello et al. (2018) reported a superior performance of NCD over *T*_2_, and that both (Siewert and Voight, 2017) and NCD outperform *T*_1_. However, in the Bitarello et al. (2018) experiments, both *T* statistics were assigned a window size of 100 informative sites on either side of the test site (*i.e*., covering 201 informative sites, which matches a physical window of approximately 10 kb), whereas NCD was assigned a window size of 3 kb. We have shown in the *Results* that after matching the window size used by each method, both the single- and trans-species variants of *T*_1_ and *T*_2_ outperform the respective single- and trans-species HKA and NCD variants, respectively. An exception is under settings with elevated mutation rates, which we further explore in the subsequent subsection *The effect of multiple tests under elevated mutation rates*. Additionally, we noticed that the powers of NCD statistics tended to be close to those of *T*_2_ when the equilibrium frequency is close to the target frequency assigned to NCD. Integrating an optimization processes into the NCD framework, however, did not improve the robustness to varying equilibrium frequencies, but instead lowered the power of the specific optimization approach that we employed (see *Supplementary Note* 6). In contrast, the performances of variants of the *T*_2_ statistics were not hindered by optimization (see *Supplementary Note* 6). Taken together, these results indicate that the model-based *T*_trans_ statistics have superior performance relative to complementary approaches for robustly identifying loci evolving under ancient trans-species balancing selection.

### The effect of multiple tests under elevated mutation rates

We observed that, regardless of whether high mutation rate was uniform across the phylogeny or was species-specific, the model-based *T* statistics reported notable increases in their proportions of false signals (Figures S8 and S9). However, because these statistics are computed at every informative site in a genomic region, regions with a higher density of informative sites due to elevated mutation rates will by chance harbor larger test scores. To examine whether this multiple testing issue can explain the inflated proportions of false signals in *T* statistics, we down-sampled the number of test sites so that the number of *T* scores computed along sequences with elevated mutation rates matched that of the original mutation rate. We considered two down-sampling approaches, which we refer to as *dense* and *sparse*. In the *dense* scenario, we sampled a specified number of test sites contiguously along the sequence, such that each pair of neighboring test sites had the same number of informative sites in common used in their computation as do those under the original mutation rate. In the *sparse* scenario, we evenly sampled the test sites along the simulated genomic region, such that informative sites distributed across the whole simulated region could be considered, but the neighboring test sites would share fewer informative sites in their computation compared to the sequences with the original mutation rate.

Indeed, either by densely (Figures S18A-C) or sparsely (Figures S18D-F) down-sampling, all *T* statistics showed substantially improved performances for sequences under uniformly elevated mutation rates (compare to Figure S8), highlighting the effect of multiple testing on inflating the proportions of false signals. For sequences with species-specific increase of mutation rates, *T* statistics also respond to down-sampling (Figure S19) in a similar fashion. Moreover, we observed that the *sparse* down-sampling scheme (Figures S18D-F and S19D-F) did not rescue the proportions of false signals as much as the *dense* scheme (Figures S18A-C and S19A-C). This performance margin echoes the fact that test sites that were sampled in the *sparse* scheme are less correlated. That is, because two neighboring test sites have fewer informative sites in common used for their calculation, even after down-sampling to match the number of sites on sequences under the original mutation rate, the set of test sites from the *sparse* scheme still has a larger effective number of tests. Therefore, multiple testing contributed to the substantially inflated mis-identification rates of *T* statistics, and when the number of tests is properly controlled, the *T* statistics (especially *T*_2,trans_) should be robust to genomic regions with high mutation rate.

Further, it is worth pointing out that this multiple-testing issue does not result from inherent statistical properties of *T* statistics, but rather the way in which windows were chosen to compute the statistics. We set the *T* statistics to perform a test on every informative site that could be centered on, and computed the likelihood ratios based on a test window with a fixed number of informative sites. As a consequence, the number of computed tests for *T* statistics will be close to the number of available informative sites, and similarly, if other statistics were computed using windows identical to *T* statistics, then their numbers of computed tests would also match the number of informative sites. When we computed single- and trans-species variants of NCD and HKA with a window size of 21 informative sites and step size of one informative site (*i.e*., making the identical number of computations on identical data as do *T* statistics) both summary statistics mimicked the behaviors of their respective model-based analogs (*i.e*., HKA to *T*_1_, and NCD to *T*_2_; Figure S20) under scenarios of high mutation rate, both when all species have high mutation rate (Figures S20A-C) and when it is specific to only species H (Figures S20D-F). These results highlight that the performance of a statistic is highly dependent on the type and number of windows it is applied with, and that the poor performance of HKA and the excellent performance of NCD under elevated mutation rate was simply due to the manner in which we computed the statistics, with physical-length windows and a fixed number of windows per simulated genomic region.

### Detecting long-term balancing selection with and without trans-species polymorphisms

Because our aim has been to circumvent potential issues surrounding trans-species polymorphisms, we only considered within-species polymorphisms and cross-species fixed differences as input data for multi-species variants of *T*_1_ and *T*_2_. Nonetheless, when conditions allow, it is possible for multi-species variants of *T*_1_ and *T*_2_ to include trans-species polymorphisms in the model (hereafter referred to as *T*_1,TSP_ and *T*_2,TSP_, respectively; see *Supplementary Note* 4), and be applied to input data with all three types of informative sites. To assess the ability of these new statistics to detect trans-species balancing selection, we examined the powers of *T*_1,TSP_ and *T*_2,TSP_ under settings in which trans-species polymorphisms are removed and in which they are included in the dataset.

When applied to the same set of simulated sequences as in previous analyses while also permitting information on trans-species polymorphisms, *T*_1,TSP_ and *T*_2,TSP_ show substantial increases in power relative to *T*_1,trans_ and *T*_2,trans_ (Figures S21), while remaining largely unaffected by balancing selection occurring after the split of the pair of species. Both statistics reached a power higher than 0.8 when selection was older than 13 million years, and *T*_2,TSP_ almost reached a power of 1.0 when the selected allele was introduced 15 million years ago. This power increase is expected due to the high probability that trans-species balancing selection would lead to trans-species polymorphisms, particularly compared to the relatively low probability expected from neutrality (*e.g*., Takahata, 1993; Leffler et al., 2013; Gao et al., 2015). Moreover, in contrast to *T*_1,trans_ and *T*_2,trans_, we observed that *T*_1,TSP_ and *T*_2,TSP_ display a gradual increase in power with increasing age of trans-species balancing selection. This pattern resembles that of single-species *T*_1_ and *T*_2_ statistics, and can partially explain why *T*_1,trans_ and *T*_2,trans_, which do not utilize information on polymorphisms established prior to species splits, maintain their powers around a constant level instead of gaining power with selection age. Moreover, when trans-species polymorphisms are absent from the data, the powers of *T*_1,TSP_ and *T*_2,TSP_, albeit respectively slightly lower than *T*_1,trans_ and *T*_2,trans_, remain similar to the *T*_1,trans_ and *T*_2,trans_ variants that do not account for trans-species polymorphisms, suggesting that the trans-species polymorphism-inclusive model can be robustly applied to data with or without trans-species polymorphisms present.

Despite this improvement in performance, however, incorporating trans-species polymorphisms in our analysis can increase the vulnerability of *T* statistics to non-adaptive processes such as mapping errors at paralogs across species and high mutation rate, both of which can lead to trans-species polymorphisms. To test the robustness of *T*_TSP_ statistics to elevated mutation rates, we applied them on sequences neutrally evolving with mutation rate five- or 20-fold higher than the original rate of *µ* = 2.5 × 10^−8^ (Figures S22A and D, respectively). In both scenarios, when trans-species polymorphisms are included in the input, *T*_TSP_ statistics falsely identified considerably more sequences as undergoing trans-species balancing selection than when such information was not provided. Moreover, higher mutation rates lead to a greater number of false signals identified by *T*_TSP_ statistics when trans-species polymorphisms are included (Figure S22). In contrast, without using trans-species polymorphisms, *T*_TSP_ statistics perform only slightly worse than *T*_trans_ statistics in mis-classifying highly mutable regions as undergoing balancing selection. On the other hand, when we applied *T*_TSP_ statistics on sequences with mutation rate 20-fold higher in species H, *T*_TSP_ statistics no longer report a greater number of false signals than do *T*_trans_ statistics (Figure S22G). This is because when the elevation in mutation rate occurred only in species H, the resulting trans-species polymorphisms would not increase as much as they did when both species H and C have high mutation rate.

To examine the effect of multiple testing on the proportion of false signals identified by *T*_TSP_ statistics, we performed down-sampling following the same procedures as described in subsection *The effect of multiple tests under elevated mutation rates*, such that regardless of whether trans-species polymorphisms are considered, the same number of test sites were sampled to infer proportions of false signals for all *T* statistics. We found that although accounting for the number of tests rescued the mis-identification rates of all *T*_trans_ and *T*_TSP_ statistics (Figure S22B, C, E, F, H, and I), the relative differences in the rates of identifying false signals between *T*_TSP_ and *T*_trans_ still persist. These results are likely due to the emphasis that *T*_TSP_ statistics place on trans-species polymorphisms, and highlight the sensitivity of *T*_TSP_ to the presence of such polymorphisms regardless of the processes that generated them. Therefore, because such trans-species polymorphisms can be generated by technical artifacts in addition to non-adaptive evolutionary processses, we recommend excluding trans-species polymorphisms while applying *T* statistics, and when such data are included, candidate regions obtained by *T*_TSP_ statistics that also harbor trans-species polymorphisms should be further validated using the framework of Gao et al. (2015), which only considers trans-species polymorphisms and is complementary to our *T* statistics.

### Applicability of trans-species methods on empirical data

In this study, we introduced a number of trans-species methods, including summary- and model-based approaches. In addition to their nuanced performances on simulated data, their applicability on empirical data also varies. In particular, two major considerations are the evolutionary relationship of the set of study species as well as the availability of sophisticated data. With respect to evolutionary relationships, divergence times between pairs of species is crucial—particularly for the model-based *T*_trans_ statistics, as their underlying models assume reciprocal monophyly. Specifically, under the Kaplan-Darden-Hudson model (Kaplan et al., 1988; Hudson and Kaplan, 1988), the tallest coalescent tree manifests at the site under long-term balancing selection, where the symmmetric population-scaled mutation rate *θ*_sel_ between the balanced alleles further controls the expected tree height. The greater the value of *θ*_sel_, the shorter time to the most recent common ancestor for the set of alleles sampled within a species (Hudson and Kaplan, 1988). To ensure monophyly at neutral sites close to the selected site, we set *θ*_sel_ = 0.05, as did DeGiorgio et al. (2014), such that the mean coalescence time within a species is approximately 12 coalescent units at the site under selection. Therefore, sister species such as humans and chimpanzees, which have a split time of approximately 12.5 coalescent units (Prado-Martinez et al., 2013), would be suitable for our current implementation of *T*_trans_ statistics. Meanwhile, for more recently diverged species such as chimpanzees and bonobos, whose inter-species divergence is approximately three coalescent units (Prüfer et al., 2012; Prado-Martinez et al., 2013), the value of *θ*_sel_ needs to be increased for *T*_trans_ to be applied. However, lowering the maximum expected tree height to such a small level would reduce the distinction between the model of balancing selection and neutrality, and likely lead to a dramatic decrease in the power of the *T*_trans_ statistics. On the other hand, the complementary summary-based HKA_trans_ and NCD_trans_ statistics can be applied regardless of the species split times.

Differences in data requirements between the summary- and model-based approaches will also influence the breadth of their application. For all methods we introduced, it is important that polymorphism data is available for at least two species, and that the reference genomes of these species can be aligned. Further, because the footprint of long-term balancing selection is small (Hudson and Kaplan, 1988), all the methods discussed here are applicable on species with draft genomes that may have relatively short scaffolds, although larger scaffolds may be important for biological interpretations. Moreover, all these methods must be applied to chromosomes that recombine, as their application hinges on the premise that only neutral variation nearby a selected site is influenced by balancing selection.

In addition to these requirements, the model-based approaches also require a recombination map, which may limit their applicability on non-model organisms. Nonetheless, we argue that adopting a uniform recombination rate should be reasonable as well, considering that recombination maps may not necessarily match across the set of study species (Smukowski and Noor, 2011). Moreover, our simulations (Figure S7) have demonstrated that dramatic changes in recombination rate and grossly mis-specified recombination maps have limited effect on the performance of *T*_trans_ statistics. Another limitation to the application of the *T*_2,trans_ statistic is that it uses derived allele frequency spectra, and therefore requires a reference genome from an outgroup species to polarize alleles as ancestral or derived. In contrast, the *T*_1,trans_ statistic (as well as HKA_trans_) is not subject to this constraint, as it does not require information on allele frequency. As a consequence, both HKA_trans_ and *T*_1,trans_ can be applied to datasets in which allele frequencies cannot be estimated well, but in which polymorphic sites may be inferred with confidence (Van Tassell et al., 2008; Cutler and Jensen, 2010; Schlötterer et al., 2014).

### Examining ancient balancing-selection shared by humans and chimpanzees

In our empirical study, we applied *T*_2,trans_ on human and chimpanzee genomic data to re-examine long-term balancing selection shared by these sister species. Without employing information from trans-species polymorphisms, we recovered well-established cases, such as the MHC and *FREM3*/*GYPE* regions, both of which were previously well-characterized with ample evidence of shared polymorphisms. We additionally reported a number of novel and relevant candidate regions with outstanding scores, such as the *GRIK1*/*CLDN17* intergenic region, the cardio health-related *SLC35F1* gene, and the ABC transporter gene *ABCA13*. Despite *SLC35F1*’s frequent association with cardiac health (Den Hoed et al., 2013; Christophersen et al., 2017; van der Ende et al., 2018), it is, in contrast, mainly expressed in brain (Uhlén et al., 2015). Meanwhile, the gene *ABCA13* encodes a lipid transporter (Prades et al., 2002), and is mainly expressed in MAPCs (Tang et al., 2010), one of the bone marrow stem cells that have high differentiation potency, and are heavily involved in wound-healing and tissue repair (Hof et al., 2007). Though both novel candidates lack empirical functional studies, they are both involved in multiple intriguing tasks, and further functional investigations could potentially substantiate their selection mechanisms.

Despite the array of trans-species statistics we presented, we chose to perform the empirical scan with *T*_2,trans_ due to the high power and robustness it shows in our simulation study, including its ability to integrate more information within a larger genomic region than summary-based approaches, thereby minimizing the noise accompanied with smaller window sizes (see *Robustness to large window sizes*). Specifically, because we observed that *T*_2,trans_ had substantially higher power than other approaches with a large window size of 100 informative sites upstream and downstream of the test site (Figure S12), we chose this window size so that *T*_2_ can make use of as much data as possible, while still maintaining reasonable power.

To infer statistical significance of each test site, we performed rigorous and extensive simulations to generate a neutral distribution for *p*-value inference. We performed *T*_2,trans_ with the exact same set of parameters used in the empirical scan, such that scores from neutral simulations were based on the empirical inter-species coalescence time, polymorphism density, and site frequency spectra as well as the same window size of 100 informative sites. Although the significant candidate regions are less than a handful, we believe the significance cutoff based on our simulations was conservative, as we did not include ubiquitous processes such as background selection that lead to overall reductions in diversity (McVicker et al., 2009; Comeron, 2014), and because the simulated datasets tended to have a higher density of informative sites due to the stringent filters applied to the empirical dataset. With these factors considered, it is sensible that the distribution of *T*_2,trans_ scores for neutral replicates is right-shifted, compared with that for the empirical data (Figure S13), further highlighting the outstanding footprints of balancing selection on our significant candidate regions—the MHC, *FREM3*/*GYPE*, and *GRIK1*/*CLDN7* loci.

### Curating and validating candidate regions

The detection of long-term balancing selection has long been hindered by numerous confounding factors, such as high mutation rate, the effects of which we have evaluated extensively (see *Results* and *Discussion*), as well as technical artifacts during sequencing and mapping, which were not accounted for in our genetic simulations. We applied stringent filters (see *Materials and Methods*) on our empirical data to remove as many regions as possible that were repetitive and that exhibited mapping issues based on available tracts. Despite that more than half of all mappable sites were removed from analyses using these filters, we still observed outstanding signals likely resulting from artifacts (*e.g*., the largest peak on chromosome 13, shown in Figure S23). To further distinguish artifacts and true signals, for high-scoring candidate regions we obtained the sequencing coverage separately averaged across human and chimpanzee samples (see *Materials and Methods*), and also performed single-species *T*_2_ scans on each of the two sister species.

For a reliably mapped genomic region, a majority of sampled individuals should harbor reasonable numbers of sequencing reads at the region, and the mean sequencing depth should not be an outlier with respect to the neighboring genomic regions or to the entire genome. Therefore, in addition to using tracts from datasets of known problematic regions (*e.g*., RepeatMasker and CRG100 alignability and mappability tracts), another approach to ensure data quality is to flag abnormalities in the numbers of sequencing reads across sampled individuals. Additionally, for regions with high data quality, it would be reassuring if footprints of long-term balancing selection also manifested separately in each species if selection occurred ancestrally and affected all species. However, this should not be a requirement, as the power to detect true signals using single-species statistics is considerably lower than with trans-species approaches.

In our results, we found that both the MHC (Figure S24) and the *FREM3*/*GYPE* (Figure S25) regions exhibited these positive features. Specifically, the peaks at these regions in single-species scans with *T*_2_ aligned well with those of the trans-species scan with *T*_2,trans_ (Figures S24A and S25A). Moreover, within either humans or chimpanzees, most of the samples reported sequencing reads, and the mean numbers of reads (*i.e*., sequencing depths) across all samples within each species at these peaks did not deviate substantially from neighboring genomic regions. Note that although the sequencing depths did fluctuate at a small region around gene *GYPB* on chromosome 4, and around *HLA-DQA* and *HLA-DQB* genes on chromosome 6, these abnormal patches either were already removed by filters prior to the scan, or did not produce high scores. Similarly, within the one Mb genomic regions covering *GRIK1*/*CLDN17* (Figure S26), *SLC35F1* (Figure S27), or *ABCA13* (Figure S28), the sequencing depths across all samples in both species are evenly distributed across the genomic regions. Meanwhile, the *T*_2_ scores of both species across these regions do not align as well as in the peaks at *HLA-DPB2* or *FREM3*/*GYPE*, and *T*_2,trans_ seems to match closer with the *T*_2_ score for chimpanzees.

In contrast, we observed potential signs of artifacts when curating other candidate regions, such as at the gene *THSD7B*, the *PGLYRP3*/*PGLYRP4* cluster, and the gene *SNTG2*. For one, although *THSD7B* (Thrombospondin Type I Domain Containing 7B) harbors an outstanding peak surrounding one of its variants (rs1469621) associated with atopic dermatitis and asthma (Figure S29; Weidinger et al., 2013), this peak region also features abnormally high numbers of reads in chimpanzee genomes (Figure S30), which often occurs at regions with problematic mapping, such as duplicated and repetitive regions. Similarly, in the case of the *PGLYRP3*/*PGLYRP4* gene cluster (Figure S31), these two genes not only encode the bactericidal peptoglycan recognition proteins (Kashyap et al., 2011, 2014), but also locate within the psoriasis-sensitive PSORS4 locus (Sun et al., 2006) with associations to several autoimmune disorders (Dziarski and Gupta, 2010; Zulfiqar et al., 2013). However, despite that innate immunity has been speculated to be a major target of balancing selection (Prugnolle et al., 2005; Ferrer-Admetlla et al., 2008; Fumagalli et al., 2009), sequencing depths around the *PGLYRP3*/*PGLYRP4* cluster in the chimpanzee genomes are also abnormally high (Figure S32), raising concerns for potential mapping issues in this region. In addition to abnormal increases of sequencing depths, an abrupt decrease in the numbers of reads may also be indicative of a region that is problematic to map, as in the case for the *SNTG2* (Syntrophin Gamma 2; Figures S33 and S34) gene on chromosome 2. As a cytoplasmic adaptor protein, γ-2 syntrophin is heavily expressed in neuronal cells (Piluso et al., 2000), and interacts with the autism-related neuroligin 3 and 4X (Yamakawa et al., 2007). However, although there are several eQTLs for its expression in testes in the peak region (Figure S33A; eQTL data from Lonsdale et al., 2013), the genomic segment immediately upstream of the peak is almost devoid of sequencing reads across the YRI samples (Figure S34B). Such regions of low coverage are suggestive of a deletion, where erroneous mapping is prone to occur, and therefore we believe that caution should be warranted before making functional interpretations at this candidate gene.

Likewise, the peak located on chromosome 13 (Figures S23 and S35A) is highly suspicious. Although this peak is a statistically-significant outlier and has some transcription factor-binding sites enriched nearby (ChIP-seq data from Ziller et al., 2013), we could not identify any coding genes within the one Mb region surrounding this peak. A closer look at the sequencing depths of this region (Figure S36) reveals an elevated plateau in depth under the peak followed by a sudden drop across the human samples (Figure S36B), suggesting that this region could harbor incorrectly mapped reads, potentially from an unfiltered duplicated region. Further, although sequencing depths in chimpanzees are evenly distributed under the peak, due to the lack of a segmental duplication filter for chimpanzee genomes, it is possible that the erroneous mapping resulted form a duplicated segment elsewhere in the genome confounded the variant calls. This suspicion is echoed when we examined the spatial distribution of allele frequencies on this region (Figures S23B and C, and Figure S37). We observed long tracks of alleles with identical moderate frequencies in the chimpanzee samples, some extending as long as almost 500 kb. Such extended tracks of identical allele frequencies are unlikely to be produced by natural processes, and further cast doubt on the validity and reliability of the signal at this region.

Lastly, it is worth noting that although these dubious signals arise on regions not included in filters such as RepeatMasker and CRG100 alignability and mappability, applying more filters does not necessarily guarantee their removal, and may discard true signals in the process. For example, when we removed all regions with copy number variation in the human genome (data obtained from Firth et al., 2009), the dubious peak on chromosome 13 was indeed successfully removed (Figure S35B). However, this additional filter halved the usable data in the process, and removed the functionally relevant and established (Prugnolle et al., 2005; Meyer et al., 2017; Leffler et al., 2017) signals at the MHC and *FREM3*/*GYPE* regions as well. We therefore recommend that comprehensive curation of candidate regions be considered when searching for genomic regions evolving under long-term balancing selection, which, in addition to extensive filters, should take both sequencing quality and potential biological mechanisms into account.

### Concluding remarks

In this study, we presented a set of methods, including both model-based and summary statistic approaches, for detecting multi-species balancing selection without dependence on the knowledge of trans-species polymorphism, and have comprehensively evaluated their performances. We have shown that all multi-species methods have specificity in detecting shared balancing selection, and have addressed how their powers could be influenced by recent demographic changes, uneven recombination rates, selection strengths and equilibrium frequencies at loci undergoing balancing selection, as well as large window sizes. We also demonstrated that our model-based approaches can be augmented to accommodate data on trans-species polymorphism to increase detection ability, but caution the use of such alterations as they can lead to false signals due to non-adaptive processes, whereas avoidance of such issues was a major impetus of this study. Application of the model-based method *T*_2,trans_ on human and chimpanzee genomic data not only recovered well-established candidates, but also revealed a number of novel putative targets that contribute to the hypothesis that pathogen defense, both in terms of adaptive immunity and of innate immunity, has been one of the prime driving forces maintaining polymorphisms across humans and chimpanzees. Lastly, we developed the software packages *MULLET* (http://www.personal.psu.edu/mxd60/mullet.html) and *MuteBaSS* (http://www.personal.psu.edu/mxd60/mutebass.html) for the respective implementation of the multi-species model-based and summary statistic approaches presented here, and we provide values for the *T*_2,trans_ test statistic and associated *p*-values from our empirical scans at http://www.personal.psu.edu/mxd60/mullet.html.

## Materials and Methods

### Simulating genetic data

We employed SLiM (Messer, 2013) to generate simulated sequence data. As recommended, we initiated each replicate simulation with a burn-in of 10*N* = 10^5^ generations, where *N* = 10^4^ is the diploid effective population size. Our simulations assumed a per-base per-generation mutation rate of *µ* = 2.5 × 10^−8^ (Nachman and Crowell, 2000), and a per-base per-generation recombination rate of *r* = 10^−8^ (Payseur and Nachman, 2000). To speed up simulations, we applied the common method of scaling parameters by a factor λ. Under this scaling, we multiplied the per-generation mutation rate, per-generation recombination rate, and per-generation selection coefficient by λ, and we divide all times in generations by λ and the diploid effective size also by λ. This scaling generates the same levels of variation expected under a simulation without scaling, except simulations run approximately λ^2^ faster, permitting an interrogation of a wider parameter space. For scenarios based on a model of constant population size or on a model of recent population expansion, we used λ = 100. For scenarios based on a model of a recent population bottleneck, we used λ = 50.

### Examining performances of two-species statistics

We simulated a demographic history analogous to that of the great apes, assuming a uniform generation time of 20 years, as did numerous prior studies (Caswell et al., 2008; Becquet and Przeworski, 2007; Langergraber et al., 2012). To comprehensively examine performances of two-species statistics, we simulated a three-species demographic history (Figure 2A) in which two sister species, analogous to humans and chimpanzees (Kumar et al., 2005), diverging *τ*_1_ = 5 × 10^6^ years ago, split with the reference species *τ*_2_ = 8 × 10^6^ years ago, analogous to gorillas (Scally et al., 2012). At the end of each simulated replicate, we sampled 50 haploid lineages from each of the sister species, and one haploid lineage from the outgroup species (species G in Figure 2A) to polarize alleles as derived or ancestral.

To examine common demographic histories that are consistent with human evolution, we simulated models with a recent population bottleneck and a recent population expansion based on parameters inferred by Lohmueller et al. (2009). Under a scenario of a recent population bottleneck, we modeled forward in time a reduction in population size from *N* = 10^4^ diploid individuals to *N_b_* = 550 diploid individuals *τ_b_* = 3.0 × 10^4^ years ago, followed by an increase in population size to *N* = 10^4^ diploid individuals *τ_e_* = 2.2 × 10^4^ years ago. Under a scenario of a recent population expansion, we modeled forward in time an increase in population size from *N* = 10^4^ diploid individuals to *N_g_* = 2 × 10^4^ diploid individuals *τ_g_* = 4.8 × 10^4^ years ago.

We generated 10^3^ replicates evolving neutrally along the three-species demographic history with constant population sizes (Figure 2A), and 400 replicates evolving neutrally under those of a recent population bottleneck (Figure 2B) or expansion (Figure 2C). For every scenario in which balancing selection was introduced, we generated 00 replicates for scenarios with selection coefficient *s* = 0.01 and dominance parameter *h* = 100, and 200 replicates for all other scenarios. For a given demographic model, we used the polymorphism and substitution data across all neutral replicate simulations to estimate the mean inter-species coalescence time 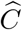, the proportions of polymorphism and substitution observed in each species, and the derived allele frequency spectra for each species (DeGiorgio et al., 2014). These quantities were provided as background genomic data for single- and trans-species variants of HKA, *T*_1_, and *T*_2_, as they require information about patterns of variation expected across the genome. For single-species variants of HKA, NCD, *T*_1_, and *T*_2_, substitutions were called as fixed differences between one ingroup species and the out-group species. However, for two-species variants HKA_trans_, NCD_trans_, *T*_1,trans_, and *T*_2,trans_, substitutions were called as fixed differences between the two sister species. In addition, for application of HKA_trans_, NCD_trans_, *T*_1,trans_, and *T*_2,trans_, we filtered out any site that was polymorphic in both sister species.

To simulate species-specific elevations in mutation rates, we employed SLiM2 (Haller and Messer, 2016, version 2.6), which allows the mutation rate to be reset, albeit globally, at a given generation. We adopted all the parameters of the three-species demographic model with constant population size (Figure 2A), and used a scaling factor of λ = 100. For each replicate, we set up two simulations, denoted as *mut* and *null*, to run in parallel using the same seed and initial model parameters. Immediately after species H and C split, all sequences in the *mut* simulation experienced a change in mutation rate, whereas the mutation parameter did not change in the *null* simulation. When simulations completed, 50 haploid lineages of species H were sampled from the *mut* simulation, whereas 50 lineages of species C and one lineage of species G were sampled from the *null* simulation. Data from these samples were then merged for downstream analyses. In this pipeline, because *mut* and *null* simulations start with the same seed and initial parameters, we can consider the evolutionary process in the two as identical until species H and C split, and that the elevation in mutation rate only occurs in species H, as we only sample lineages from species H in the *mut* simulation.

We considered three scenarios in which the mutation rate was increased to 5*µ*, 10*µ*, or 20*µ*, with *µ* being the initial mutation rate of 2.5 × 10^−8^ per site per generation, which was used throughout the *null* simulations. We also simulated one additional scenario where neither the *mut* nor the *null* simulation reset their mutation rate and instead remained constant at rate *µ*. We applied the same sampling and parsing procedures to this set of simulations, and used these simulation replicates to generate a null distribution to obtain false positive rates. We simulated 400 replicates for each of these four scenarios, and performed the same analysis procedures as in other simulation settings. We supplied single- and trans-species variants of HKA and *T*_1_ with proportions of polymorphisms and substitutions inferred from the concatenated sequences from the constant mutation rate scenario, from which we also derived the inter-species coalescence time and allele frequency spectra for the application of *T*_2_ variants. All summary statistics were applied with windows of size one kb, and model-based *T* statistics with 10 informative sites flanking each side of the test site.

### Assessing power of *K*-species statistics

To assess the ability of *K*-species (*K* = 2, 3, or 4) variants of NCD, HKA, *T*_1_, and *T*_2_ to detect and localize sites undergoing ancient balancing selection, we included two additional species onto the three-species demographic history in Figure 2A, with the fourth and fifth species diverging *τ*_3_ = 12 × 10^6^ and *τ*_4_ = 17 × 10^6^ years ago, respectively, analogous to that of orangutans (Auton et al., 2012) and gibbons (Carbone et al., 2014) (respectively denoted by species O and B in Figure 5A). We simulated 50 kb-long sequences evolving along the tree, with a uniform recombination rate of *r* = 10^−8^ per site per generation, and a mutation rate of *µ* = 2.5×10^−8^ per site per generation across all species, consistent with our previous simulations. We also assumed a constant population size of *N* = 10^4^ diploids across the entire phylogeny. Fifty haploid lineages were sampled each from species H, C, G, and O in the present, and one lineage was sampled from species B. Derived alleles were called based on a single sampled lineage from the nearest outgroup species to the set of *K* species examined. That is, we used one sampled lineage from species C to polarize alleles in species H for one-species experiments, species G to polarize alleles in species H and C for two-species experiments, species O to polarize alleles in species H, C, and G for three-species experiments, and species B to polarize alleles in species H, C, G, and O for four-species experiments.

We generated 400 neutral replicates under the five-species demographic history, and 200 replicates for each scenario featuring alleles under balancing selection. We parsed each replicate such that the informative sites include only intra-species polymorphisms and inter-species substitutions that agree with the phylogeny. We provided *K*-species HKA variants with the proportion of each type of informative site computed across the entire set of neutral replicates. We provided *K*-species *T*_1_ and *T*_2_ variants with polymorphism and substitution configurations and site frequency spectra computed across the entire set of neutral replicates, in addition to the inter-species coalescence times calculated from known simulation parameters. We applied all *K*-species variants of HKA and NCD with sliding windows of length one kb with a step size of 500 nucleotides to advance the window. To match the approximate quantity of data utilized by each summary statistic, we provided *K*-species variants of *T*_1_ and *T*_2_ with 10 informative sites on either side of the test site.

### Empirical data analyses

We used allele frequency data of all 10 unrelated western chimpanzee individuals from the PanMap Project (Auton et al., 2012), as well as 108 African human individuals of the Yoruban (YRI) population from the 1000 Genomes Project (The 1000 Genomes Project Consortium, 2015). The data for the two species were originally mapped to the chimpanzee panTro2 (March 2006, Washington University Build 2 Version 1) and the human hg19 (February 2009, GRCh37 Genome Reference Consortium Human Reference 37) reference genomes, respectively. We mapped the chimpanzee variant call data to hg19, and used the sequence information from the gorilla gorGor3 reference genome (Kent et al., 2002, downloaded from UCSC Table Browser) to polarize alleles as ancestral or derived. Aligned sites that are not included in the variant call datasets were assumed to be monomorphic for the reference allele, and only bi-allelic single-nucleotide sites segregating across lineages from all three species were considered. Moreover, we only considered genomic regions mappable among all three species, and regions of human reference genome hg19 that did not uniquely map to chimpanzee reference genome panTro2 were also removed.

To circumvent potential mapping issues from paralogs, we performed one-tailed tests on both chimpanzee and human data for Hardy-Weinberg equilibrium as described by Wigginton et al. (2005), and discarded sites with excessive heterozygosity in each dataset as determined by *p*-values less than 10^*−*^4. For the one-tailed Hardy-Weinberg test in humans, we used human genotype data from all individuals in the 1000 Genomes Project, as mapping issues would manifest across all populations, and the larger sample would increase power to reject Hardy-Weinberg equilibrium. Moreover, this pooling of individuals is not affected by the Wahlund effect (Wahlund, 1928), as we specifically performed a one-tailed test to uncover sites with excess heterozygotes rather than also testing for excess homozygotes. Further, we discarded genomic regions with 100-mer mappabilities (*i.e*., CRG scores computed by Derrien et al., 2012) less than 1.0. We also removed all variants that intersected segmental duplications annotated in hg19 or simple repeats annotated in panTro2, as well as all repetitive regions in both hg19 and panTro2 suggested by repeatMasker (all obtained from UCSC Table Browser; Kent et al., 2002). Matching the filtered data, we adopted the inferred recombination map of human hg19 reference genome (The International HapMap Consortium, 2007), and removed the chromosomal regions not covered by the map. We then examined the proportions of polymorphism and substitution, derived allele frequency spectra, and estimated human-chimpanzee inter-species coalescence time (Prado-Martinez et al., 2013) for consistency with our expectations.

To assign *p*-values to test sites in our empirical scan, we employed the coalescent simulator *ms* (Hudson, 2002) and generated 5 × 10^7^ independent replicates of 25 kb-long sequences. We simulated a three-species demographic history, with parameters for the YRI human population adopted from Terhorst et al. (2017), and from Prado-Martinez et al. (2013) for chimpanzees and gorillas. Specifically, for temporal changes in great ape population sizes, we used the parameters estimated by PSMC in Prado-Martinez et al. (2013). For humans, we adopted the SMC++ estimates of Terhorst et al. (2017). For the split times between humans and chimpanzees, and between humans and gorillas, we adopted the ILS CoalHMM estimates reported by Prado-Martinez et al. (2013). For all simulated sequences, we assumed a generation time of 20 years, a uniform mutation rate of 1.25 × 10^−8^ per site per generation and a uniform recombination rate of 10^−8^ per site per generation (as used by Terhorst et al., 2017). For each sequence, we applied *T*_2,trans_ on the first 201 informative sites to compute a single score at the 101th informative for each simulated replicate, with the same parameters adopted in the empirical scan, including inter-species coalescence time, genome-wide proportions of polymorphisms and substitutions, as well as genome-wide estimates of the site frequency spectra of both species.

To further interrogate the footprints uncovered in candidate regions, we re-examined the sequencing quality in both humans and chimpanzees at these regions. We obtained the BAM files of mapped sequencing reads for each human and chimpanzee individual included in this study, and generated whole-genome maps of sequencing depth across individuals. For each genomic region, we computed the mean sequencing depth across individuals from the same species, and flagged regions with more than half of the sampled individuals devoid of sequencing reads. Furthermore, to investigate whether copy number variation could have contributed to false signals due to technical artifacts, we took advantage of the human copy number variation map provided by the DECIPHER project (Firth et al., 2009), and performed another scan with *T*_2,trans_ applied to data with regions containing copy number variation further filtered. Moreover, we also separately applied the single-species *T*_2_ statistic (DeGiorgio et al., 2014) on human and chimpanzee data to examine whether there was concordance with shared signals identified from our *T*_2,trans_ scan. Specifically, to ensure the data used in the scans within each species were comparable, we only considered sites that were mappable across hg19, panTro2, and gorGor3 reference genomes, and that have passed all the aforementioned filters. For application to human (chimpanzee) data, we applied *T*_2_ on sites that were either polymorphic within human (chimpanzee) or substitutions with respect to gorGor3.

## Acknowledgments

We thank Michelle S. Kim for her contribution in testing the draft software for two-species versions of the *T*_trans_ statistics, Javier Prado-Martinez for providing PSMC estimates of great ape demographic history estimates from Prado-Martinez et al. (2013), and Jonathan Terhorst for providing SMC++ human demographic history estimates from Terhorst et al. (2017). We also thank two anonymous reviewers for their constructive feedback that helped strengthen this manuscript. This work was funded by National Institutes of Health grant R35GM128590, by the Alfred P. Sloan Foundation, and by Pennsylvania State University startup funds. Portions of this research were conducted with Advanced CyberInfrastructure computational resources provided by the Institute for CyberScience at Pennsylvania State University. This study makes use of the copy number variation data generated by the DECIPHER community. A full list of centers who contributed to the generation of the data is available from http://decipher.sanger.ac.uk and via email from. Funding for the project was provided by the Wellcome Trust.

